# L-Serine and Palmitoyl CoA control *Mycobacterial* infection by tweaking protective immune response: A potential host directed therapy for tuberculosis

**DOI:** 10.64898/2026.08.20.745965

**Authors:** Niharika Sharma, Rahul Sharma, Abhishek Kumar, Lalit Kumar Singh, Anjaneya Ayanur, Vijay Hadda, Amit Kumar Singh, Hridayesh Prakash

**Affiliations:** Amity Centre for Translational Research, Amity University, Noida 201313, India; ICMR-National JALMA Institute for Leprosy and Other Mycobacterial Diseases, Agra 282004, Uttar Pradesh, India; CSIR-Indian Institute of Toxicology Research (CSIR-IITR), Lucknow 226008, Uttar Pradesh, India; Department of Pulmonary, Critical Care & Sleep Medicine, All India Institute of Medical Sciences, New Delhi 110029, India; Academy of Scientific and Innovative Research (AcSIR), Ghaziabad, Uttar Pradesh 201002, India

**Keywords:** L-Serine, Sphingolipids, Host-directed therapy, Tuberculosis, Rifampicin, Multidrug-resistant TB (MDR-TB), Nitric oxide, Adjunctive therapy, Immunomodulation

## Abstract

Host-directed therapies (HDTs) have acquired paramount importance for management of tuberculosis (TB). L-Serine is an important metabolite and immunomodulatory biomolecule with promising role in managing infections, and autoimmune diseases. However, the role of L-Serine as host directed therapeutic against *Mycobacterium tuberculosis* (*M.tb*) remains unexplored. Therefore, we adopted *de novo*-based approach and employed L-Serine and palmitoyl CoA precursors for enhancing sphingolipid in host and accessed their anti-tubercular potential. In this study, we investigated whether L-Serine could modulate the antibiotics efficacy against *M.tb*. L-Serine particularly in combination with palmitic acid, rifampicin and isoniazid showed enhanced intracellular bacterial clearance in a dose- and time-dependent manner in murine and human macrophages. This synergistic effect was accompanied by increased nitric oxide production and modulation of the host immune response. We identified elevated levels of pro-inflammatory cytokine TNF-α and reduced anti-inflammatory IL-10 expression. Furthermore, L-Serine supplementation demonstrated antimicrobial activity in isolated primary CD14⁺ monocytes from TB patients. Similarly, the metabolic supplementation of L-Serine in combination with isoniazid, rifampicin, and palmitic acid significantly reduced bacterial burden in the lungs and spleen, while improving tissue architecture in murine infection model. Our observations suggest that L-Serine and palmitic acid in combination with isoniazid and rifampicin contributes to the observed therapeutic effects. Collectively, this study concludes that L-Serine and palmitic acid acts as a promising host-directed therapeutic adjunct, which enhances antimicrobial immunity and potentiating antibiotic efficacy, providing a potential strategy for improving tuberculosis treatment outcomes.

**Importance:** TB patients are poor in their sphingolipids metabolites which are associated with their poor immunity against TB, therefore a targeted and safe approach is essential for augmenting sphingolipid levels in host for improving disease outcome. Here we have developed a safe and effective sphingolipid mimetic (L-Serine and Palmitoyl CoA) as strong adjunct for anti-tubercular drugs. We believe inclusion of sphingolipid mimetic in anti-tubercular regimen would improve drug response in MDR-TB patients by enhancing the therapeutic effect of standard anti-TB medications, potentially leading to more effective bacterial clearance. We also believe that our mimetic would shorten the treatment regimen by enhancing the effectiveness of the drugs, potentially leading to faster resolution of infection. This study delivers L-Serine / Palmitic Acid as new adjunct to anti-tubercular drugs. We have already patented this formulation and now heading for the randomized controlled trial in active PTB and MDR TB patients.

## Introduction

Tuberculosis (TB) is the leading infectious disease, with 10.7 million incidences and claiming approximately 1.25 million lives worldwide [1]. The global advancement towards the WHO End TB targets has been recalibrated to 2035. The complexity of TB prevention and treatment has been amplified due to the divergent multi-drug resistant (MDR) TB. The emergence of MDR *M.tb* strain further complicated the TB management. These challenges highlight the urgent need for innovative new treatments or adjunctive strategies. Host-directed therapies (HDTs) are emerging as a paradigm shift for effective tuberculosis management. HDT mainly targets the host immunometabolic pathways to enhance antibiotic efficacy and mitigate immunopathology. It also helps in reducing the emergence of drug resistance without implementing the selective pressure on the pathogen [2–4]. Numerous HDT candidates, including metformin, statins, and vitamin D, have demonstrated adjunctive benefits in both experimental and clinical settings against TB. Nonetheless, the metabolic pathways crucial to TB pathogenesis targeted by the HDT are limited.

The adjunctive strategies have the potential to modulate host immune and metabolic pathways to enhance bacterial clearance, limit immunopathology, and shorten treatment duration [5]. In this context, our previous work has demonstrated that modulation of host sphingolipids or bioactive amphipathic lipids can regulate immune function and enhance host defence. This host defence is mediated by promoting leukocyte trafficking, macrophage activation, mycobacterial phagocytosis, granuloma integrity, and the resolution of inflammation [6]. The altered sphingolipid metabolism has been implicated in chronic lung diseases, including chronic obstructive pulmonary disease, asthma and osteoarticular TB. Moreover, dysregulated and reduced pulmonary sphingolipid levels are associated with increased disease severity and impaired host immunity [7, 8]. Among these metabolites, sphingosine-1-phosphate (S1P) enhances antimycobacterial activity by promoting macrophage differentiation, nitric oxide production, phagosomal maturation, and restriction of intracellular *M.tb* survival [9–11]. Clinical studies further demonstrate significant depletion of sphingolipids, particularly S1P in active TB patients [13]. Although exogenous S1P supplementation reduced the mycobacterial burden in experimental mouse TB models [9]. However, S1P induces potential allergic and autoimmune responses, that limit the therapeutic applications [12].

To address this limitation, we employed L-Serine (LS) in combination with palmitic acid (PA) based on novel adjunctive strategy, which required to enhance the endogenous oxy-sphingolipid biosynthesis through the *de novo* pathway. This metabolically driven approach offers a potentially safer means of restoring protective sphingolipid pools. In parallel, this methodology simultaneously enhances the efficacy of conventional anti-tubercular therapy [13]. We demonstrate that LS exhibits modest intrinsic antimycobacterial activity and displays strong synergistic interactions (FICI ≤ 0.5) with rifampicin and moxifloxacin against MDR *M. tuberculosis*. In primary human CD14⁺ monocytes and murine macrophages, LS and PA in combination with isoniazid and rifampicin significantly enhanced intracellular mycobacterial killing and nitric oxide production. Although, the high dose of LS and PA modestly reduced pulmonary bacterial burden when administered alone in a murine pulmonary TB model. Furthermore, the adjunctive treatment with isoniazid and rifampicin achieved almost complete bacterial sterilization, complete spleen sterilization, restoration of lung architecture, and a favorable shift in the TNF-α/IL-10 balance. Together, these findings identify LS as safe, metabolically driven host-directed therapeutic adjuncts. This metabolite adjunctive therapy has the potential to improve treatment outcomes in both drug-sensitive and drug-resistant tuberculosis.

## Materials and Methods

### Reagents

We purchased most of the reagents and chemicals used in this study from the manufacturer Sigma-Aldrich (UK) e.g., L-Serine (cat. no. S-4500), palmitic acid (cat. no. P5585), glycerol, polysorbate 80, penicillin - streptomycin (cat. no. 15140-122), Lipopolysaccharide (L6511), Sulphanilamide (cat. no. S9251), N-(naphthyl ethylenediamine dihydrochloride) (cat. no. 684147), Formaldehyde (cat. no. 252549), resazurin dye, Oil Red O solution (cat. no. O1391), cell culture media RPMI-1640 (cat. no. SH30027001), DMEM, FBS (cat. no. 10270-106), Ficoll-Hypaque (cat. no. 10771) and human AB serum. The bacterial culture media e.g., Middlebrook 7H9, 7H10, ADC, OADC supplements were purchased from BD Difco (USA). We procured the recombinant cytokines TNF-α, IL-6 and IL-10 from eBioscience (San Diego, CA, USA). Microbeads and cell separation columns for mouse CD11b⁺ and human CD14⁺ monocytes were purchased from Miltenyi Biotec (Germany). We procured ELISA kits for TNF-α, IFN-γ, IL-6 and IL-10 from R&D Systems (Germany/USA). For gene expression analysis RNA isolation kits (RNeasy Mini kit, Qiagen - cat. no. 74104), SYBR Green Master Mix (BioRad - cat. no. 1725271), and PCR primers were bought from Eurofins Genomics, Bengaluru. We purchased isoniazid (cat. no. - I3377), rifampicin (cat. no. - TA9H97BAE8A6), moxifloxacin (cat. no. - AMBH2D6FDB0D), sodium nitroprusside (SNP), and cobalt chloride (CoCl₂) from Merck (USA). We procured the plasticwares and consumables from Axygen-Genaxy (USA). *Mycobacterium tuberculosis* H_37_Rv and clinical multidrug-resistant *M. tuberculosis* isolates were maintained at ICMR-NJIL&OMD, Agra.

### Resazurin Microtiter Assay (REMA)

We determined the antimycobacterial activity of LS, both individually and in combination with isoniazid, rifampicin and moxifloxacin using the resazurin microtiter assay (REMA) with slight modifications [14]. We grew the *M.tb* H_37_Rv and a clinical MDR *M.tb* strain (SN/MC-90) in Middlebrook 7H9 supplemented with 10% OADC, 0.2% glycerol, and 0.05% Tween-80 at 37°C until mid-log phase (OD_600_ 0.4–0.6). We diluted the cultures at approximate concentration of 1 × 10⁵ CFU/mL, and 100 µL was dispensed into sterile flat-bottom 96-well plates. Furthermore, we added LS, rifampicin, isoniazid, moxifloxacin, and their combinations in two-fold serial dilutions. These dilutions were prepared in 7H9 broth and a volume of 100 µL was transferred per well. We used bacteria only as growth controls and medium only as sterility controls. Then we sealed the plates with parafilm and incubated at 37°C in BSL-3 incubator for 7 days. Subsequently, 30 µL of 0.01% resazurin solution was added to each well and incubated for an additional 24–48 h. The blue wells indicated the growth inhibition, while wells that turned colour pink indicated bacterial growth. The minimum inhibitory concentration (MIC) was defined as the lowest concentration preventing colour change. Drug interactions were evaluated using the fractional inhibitory concentration index (FICI): FICI = (MIC_A in combination / MIC_A alone) + (MIC_B in combination / MIC_B alone). Synergy was defined as FICI ≤ 0.5; additive effect as 0.5 < FICI ≤ 1; indifference as 1 < FICI ≤ 4; and antagonism as FICI > 4. All experiments were performed in triplicate and repeated independently.

## Cell Culture

RAW264.7 murine macrophages and THP-1 human monocytic cells were obtained from the National Centre for Cell Science (NCCS), Pune, India. RAW cells were maintained in RPMI-1640 and THP-1 human monocytes in RPMI-1640 supplemented with 10% FBS and 1% penicillin–streptomycin at 37°C and grown in humidified incubator supplemented with 5% CO₂. Brewer thioglycolate 4% medium (1 mL) was injected into C57BL/6J mice intraperitoneally to induce macrophage recruitment [15]. Peritoneal exudate cells were collected after 72 hours by lavage in phosphate-buffered saline (PBS). Cells were centrifuged at 400 × g for 8 minutes at 4°C, and the resulting pellet was resuspended in serum-free RPMI-1640 medium. Macrophages CD11b⁺ were enriched using magnetic-activated cell sorting (MACS) according to manufacturer’s guidelines (Miltenyi Biotec, Germany). Macrophages were enriched and cultured overnight before the experimental treatment. Bone marrow cells were collected from the femurs and tibiae of C57BL/6J mice with slight modifications [16]. Following the lysis of red blood cells, the cells were incubated and cultured in complete DMEM supplemented with 10% FBS and GM-CSF (20 ng/mL) for 5 days to facilitate macrophage differentiation. Adherent macrophages were collected and utilized in subsequent experiments. Human CD14⁺ monocytes were isolated from peripheral blood (10 mL), was obtained from healthy controls (n = 10) and pulmonary TB patients, consisting of drug-sensitive (n = 10) and MDR cases (n = 10), at AIIMS, New Delhi. PBMCs were isolated by Ficoll density gradient centrifugation. Those samples were not considered into final data analysis, where the monocytes density was very low, especially in PTB or MDR samples and were excluded from experimental analysis. Human CD14⁺ monocytes were isolated and purified utilizing magnetic microbeads (Miltenyi Biotec). Cell purity (>95%) was confirmed by flow cytometry prior to the subsequent experiments.

### Cellular Infection

Macrophages were plated at 1 × 10⁵ cells/well and infected with *M. tuberculosis* H_37_Rv at an MOI of 10:1 for 3 h. After washing to eliminate extracellular bacteria, cells were treated with LS, palmitoyl-CoA/palmitic acid, standard anti-tubercular drugs (isoniazid and rifampicin), or combinations thereof. At 48 and 72 h post-treatment, cells were lysed using 0.1% Triton X-100, and lysates were diluted and plated on 7H11 agar [17]. The colony count was determined after 21 days, with the intracellular bacterial burden was represented as colony-forming units (CFU) per well. For the evaluation of innate immune response, NO production was measured using the Griess assay and cytokine levels (TNF-α, IL-10, IL-6) were determined in culture supernatants using ELISA following the manufacturer’s guidelines [18].

### Drug Preparation and Treatment Regimen

LS dose (LS^LD^: 600 mg/kg and LS^HD^: 1200 mg/kg) was dissolved in sterile PBS immediately before administration. Palmitic acid was dissolved in absolute ethanol and supplemented with 10% fatty acid-free bovine serum albumin (BSA) in PBS by heating to 70 °C with constant stirring until fully solubilized, as previously described [19, 20]. The final complex was filter sterilized through a 0.22 μm syringe filter, and aliquots were stored at −20 °C. Fresh working solutions were prepared on the dosing day. Based on literature review and pilot toxicity studies, PA doses 150 mg/kg and 300 mg/kg body weight were selected for the *in-vivo* experiments [21–23]. Treatment started at fourth week post-infection and continued for the next eighth week.

### Animal challenge study

Female BALB/c mice age six- to eight-weeks, were infected with *M. tuberculosis* H_37_Rv via aerosol challenge model (∼100–120 CFU/lung) utilizing a Glass Col inhalation system in BSL-3 facility [24]. The treatment began from 4^th^ week after infection and continued till 8^th^ week. Experimental groups were administered LS, PA and their combinations with isoniazid 25 mg/kg + rifampicin 10 mg/kg. Mice were euthanized for the analysis of mycobacterial growth, and their lungs and brains were aseptically extracted, weighed, and homogenized. Serial dilutions were plated on 7H11 agar and incubated for 3–4 weeks before the CFU enumeration. For histopathological assessment lung and spleen tissues were first fixed in formalin and then embedded in paraffin. Thin sections at 5 µm were made and stained with hematoxylin and eosin (H&E) according to standard protocol. A predefined semi-quantitative descriptive grading system was utilized for histopathological evaluation to determine the severity of tissue pathology. Lung and spleen sections were analyzed for granuloma like lesion development and multifocal collection of mononuclear cell aggregates. We analyzed the infiltration of inflammatory cells (including polymorphonuclear infiltration and thickening of alveolar septa), caseous necrosis, atelectasis and/or emphysematous alterations, and along with interstitial fibrosis. Every parameter was graded descriptively as follows: – = normal (no detectable lesion); +1 = mild focal involvement with minimal architectural distortion; +2 = moderate multifocal involvement with evident structural alteration; +3 = severe or extensive involvement with marked architectural disruption. Spleen sections were assessed for hyperplasia or proliferation of the white pulp and reduction of red pulp area. Furthermore, we analyzed trabecular fibrosis with architectural distortion, and reduced hemosiderin accumulation utilizing the same descriptive grading scale (–, +1, +2, +3). All histopathological evaluations were conducted independently in a blinded fashion by an experienced pathologist who was unaware of the experimental group allocation at the time of scoring. For analyzing immune response for lung homogenates were prepared in PBS containing 0.1% Triton X-100 and protease inhibitors. Following centrifugation at 13,000 rpm for 10 minutes at 4°C, supernatants were collected and total protein concentration was determined using a BCA assay (Thermo Scientific). Cytokine levels (TNF-α and IL-10) were measured using Duo Set® ELISA Development Kits (R&D Systems, Minneapolis, MN; DY410 and DY417) according to the manufacturer’s instructions. ELISA plate readings were taken at 450 nm with a reference at 570 nm using a microplate reader. Cytokine levels were normalized to total protein and expressed as pg/mg protein [18, 24].

### Induction of Foamy Macrophages by Lipid Treatment

For the induction of foam cell formation, adhered macrophages were stimulated with different lipid molecules like oxidized LDL (oxyLDL) (5 µg/ml), lipopolysaccharide (LPS) (100 ng/ml), palmitic acid (100 mM), naïve LDL (100 µg/ml) and oleic acid (200 µM) with incubation for 24 hours 5% CO_2_ incubator at 37^°^ C to induce foamy macrophage phenotype and lipid droplet accumulation in macrophages [25–28]. Untreated macrophages were served as experimental control. Oil Red staining was performed to confirm the foamy macrophage phenotype [29]. Following the induction of foamy macrophage by different lipid molecules, cells were subjected to metabolic intervention with L-Serine. Macrophages which induced foamy phenotype were treated with L-Serine (100 µM) for 24 hours under standard culture conditions. Effects of L-Serine on foamy macrophages were confirmed by Oil Red staining.

### Oil Red O Staining

Oil Red O staining was used to demonstrate the accumulation of lipid accumulated macrophages. The working solution of Oil Red stain must be prepared fresh and should be utilized within two hours. Following two washes PBS, then cells were fixed in 10% formalin for 30 minutes, followed by washing with Milli-Q water twice [30, 31] then washed for 5 minutes in 60% isopropanol to facilitate neutral lipid staining easier. The cells were treated with working Oil Red solution for 10 minutes and then washed 3-4 times with Milli-Q water and observed under the phase contrast microscope.

## Results

### L-Serine Augments the Efficacy of First- and Second-Line Anti-Tubercular Drugs

L-Serine (LS) is the widely accepted dietary supplement and demonstrated as adjunctive therapeutic against the *Acinetobacter* and *Streptococcus*. In our initial experiments, we aimed to examine the effects of rifampicin and moxifloxacin along with L-Serine as adjunctive treatments. We found that LS inhibits the growth of *M.tb* H_37_Rv in a dose-dependent manner. We performed the resazurin microtiter assay (REMA), which determines the reduced bacterial metabolic activity as indicated by colour change from pink to blue. In this experiment, both drug combinations demonstrated synergistic interactions with Fractional Inhibitory Concentration Index (FICI) values of ≤0.5 **(Supplementary Table 1).** Rifampicin demonstrated strong antimycobacterial efficacy with a minimum inhibitory concentration (MIC) of ∼ 0.07 µM. Conversely, LS displayed relatively weak intrinsic activity with a MIC of 50 mM **(Supplementary Figure 1A)**. Furthermore, the checkerboard assays showed that LS at concentrations ≥0.78 mM enhance the efficacy of sub-inhibitory concentrations of rifampicin (0.01–0.04 µM). These results indicate improved bacterial lethality as shown by significant blue colour shift **(Supplementary Figure 1B)**. The FICI values were observed in between a range from 0.16 to 0.52, indicating significant synergistic activity **(Supplementary Figure 1C)**. Additionally, we also examined the LS synergy with moxifloxacin against the MDR *M. tuberculosis* clinical strain SN/MC-90. LS inhibited bacterial growth solely at approximately 400 mM. Meanwhile, the combination of moxifloxacin and rifampicin demonstrated dose-dependent inhibition with MICs of about 0.62 µM and ≥2.5 µM, respectively. The above observations align perfectly with the drug-resistant phenotype **(Supplementary Figure 1D, 1E)**. Significantly, LS potentiated the effect of moxifloxacin’s antimycobacterial activity, reducing its effective inhibitory concentration to 0.31 µM. Similarly, LS improved the rifampicin induced inhibition at concentration of ≥1.5 µM **(Supplementary Figure 1E)**.

Furthermore, we examined the combinatorial impact of LS on macrophages infected with *M.tb.* Murine macrophages infected with *M.tb* were treated with LS (50 mM/ 100 mM) along with isoniazid (H), rifampicin (R) and palmitoyl CoA (PCoA) or various combinations. The intracellular bacterial load was quantified by serial dilution and colony-forming unit (CFU) enumeration was performed at 48 and 72 hours after treatment. The LS was examined at two different concentrations at 50 mM (LS^LD^:low dose) and 100 mM (LS^HD^:high dose). The low dose LS treatment demonstrated the diminished intracellular survival of bacteria, which were aligned with the in-vitro results. Moreover, the combination of LS with isoniazid and rifampicin significantly improved the bacterial clearance comparatively at both 48 and 72 hours post-infection. Meanwhile, palmitoyl CoA alone showed no notable impact. The combination of palmitoyl CoA with LS enhanced the bacterial clearance at 72 hours, indicating a time dependent synergistic effect (**Figure 1A, B; *p* < 0.05)**. Further, we estimated the impact of high dose LS on intracellular bacterial survival. The combination of LS^HD^ with isoniazid and rifampicin resulted in a strong antimycobacterial effect, decreasing intracellular bacterial load by 50% compared to untreated controls (**Figure 1C, D)**. The experimental evidence suggests that the high dose of LS with isoniazid and rifampicin led to the most significant decrease in intracellular bacteria and inclusion of palmitoyl CoA further improved the bacterial elimination. The translational significance of these results was additionally assessed using THP-1-derived human macrophages infected with *M.tb* H_37_Rv. The low dose treatment of LS combined with isoniazid and rifampicin significantly decreased the intracellular bacterial burden compared with untreated controls at 48 and 72 hours. Palmitoyl CoA had negligible effect on the survival of intracellular bacteria (**Figure 1E, F; *p* < 0.05)**. We identified that LS and palmitoyl CoA did not exhibit antimycobacterial activity when tested against the MDR strain (SN/MC-90), individually or in combination. Although, the intracellular survival was diminished with the combination of LS and palmitoyl CoA, it was not significant at 48 and 72 hours post infection **(Supplementary Figure 2A, B)**. The intracellular survival of *M.tb* in macrophages derived from primary human CD14⁺ monocytes were not reduced significantly when combination with isoniazid and rifampicin at 48 and 72 hours post infection. **(Supplementary Figure 2C, D)**.

**Figure 1.**
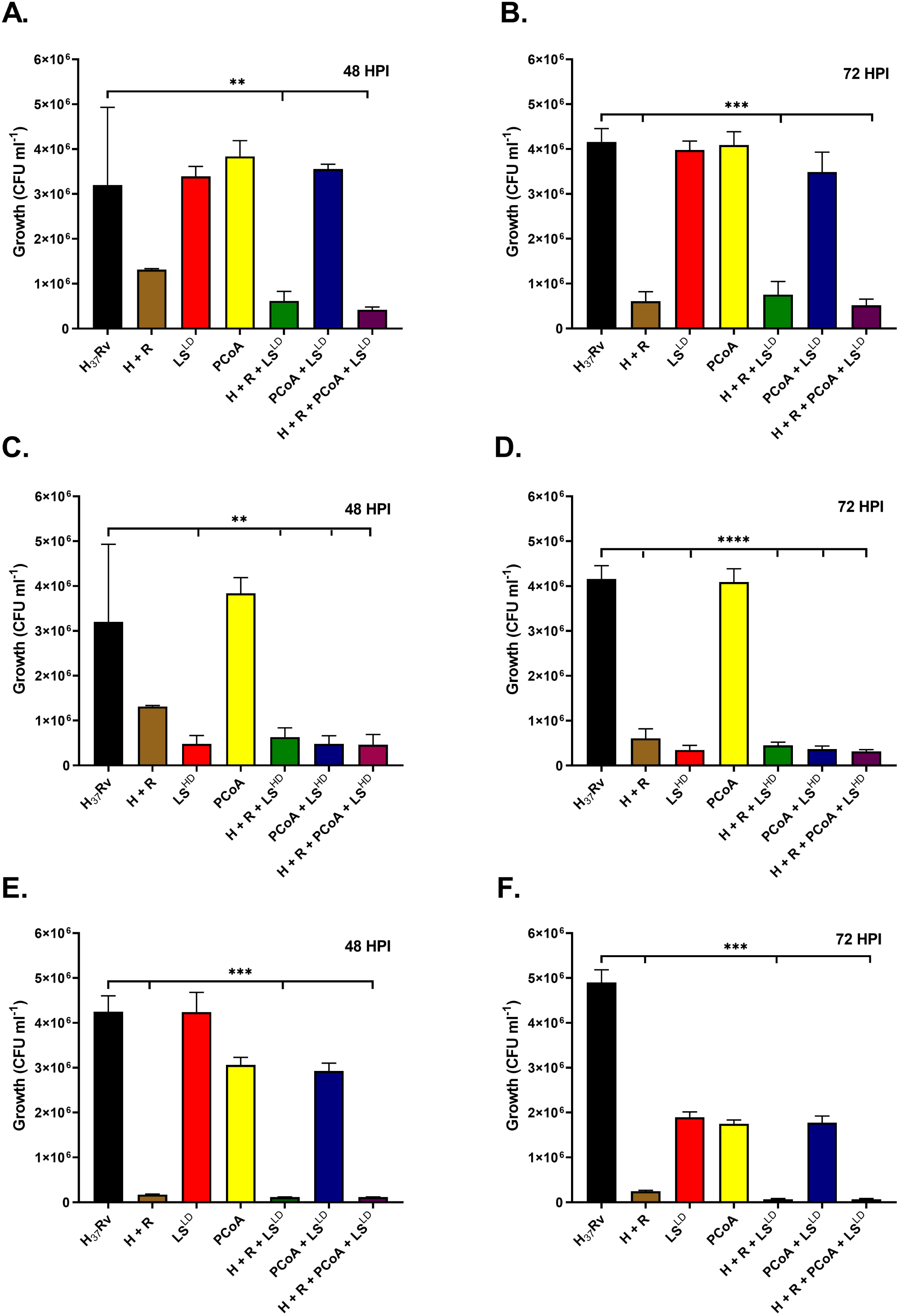
LS decreases the intracellular survival of *Mycobacterium tuberculosis* in both human and murine macrophages. L-Serine (LS^LD^: 50 µM, LS^HD^: 100 µM) was administered in combination with Isoniazid (H – 2 µM), Rifampicin (R – 2.4 µM), and Palmitoyl CoA (PCoA, 50µM) to treat *M. tuberculosis* (*M.tb*, H_37_Rv)-infected RAW264.7 macrophages. Colony-forming units (CFUs) were enumerated **(A)** 48 and **(B)** 72 hours post infection (HPI), and serial dilution was used to determine the intracellular bacterial burden. The LS was tested at a low dosage of 50 mM (LS low dose: LS^LD^). LS 100 mM (LS^HD^:high dose), PCoA, H+R and their combinations were administered in *M.tb* infected RAW264.7 macrophages. CFU enumerated **(C)** 48 and **(D)** 72 hours post infection LS 50 mM (LS^LD^:low dose), PCoA, H+R and their combinations were administered in *M.tb* infected THP-1 macrophages. CFU were enumerated **(E)** 48 and **(F)** 72 hours post infection. Data are presented as mean ± SD from three independent experiments. Statistical significance was determined by one-way ANOVA followed by the appropriate post hoc test ** p < 0.05, *** p < 0.01, and **** p < 0.001.

### L-Serine Promotes Innate Immune Responses and Activates Sphingolipid Biosynthetic Pathways in Monocytes

Nitric oxide (NO) serves as a key host defence mechanism that drives the bacteria into a dormant, drug tolerant state. That’s why estimation of NO levels is essential in understanding the *M.tb* pathogenesis. We performed the NO level estimation during LS, LPS and cobalt chloride treatments. We observed enhanced NO production in naïve RAW264.7 macrophages during LS treatment and additionally found the significantly amplified NO generation triggered by LPS and sodium nitroprusside (SNP) (**Figure 2A-B)**. The NO levels were significantly elevated also by the hypoxia mimetic cobalt chloride (**Figure 2C)**. These results suggest that LS increases nitric oxide- and hypoxia-associated signalling pathways. All these pathways play an essential role in intracellular regulation and elimination of *M.tb*.

**Figure 2.**
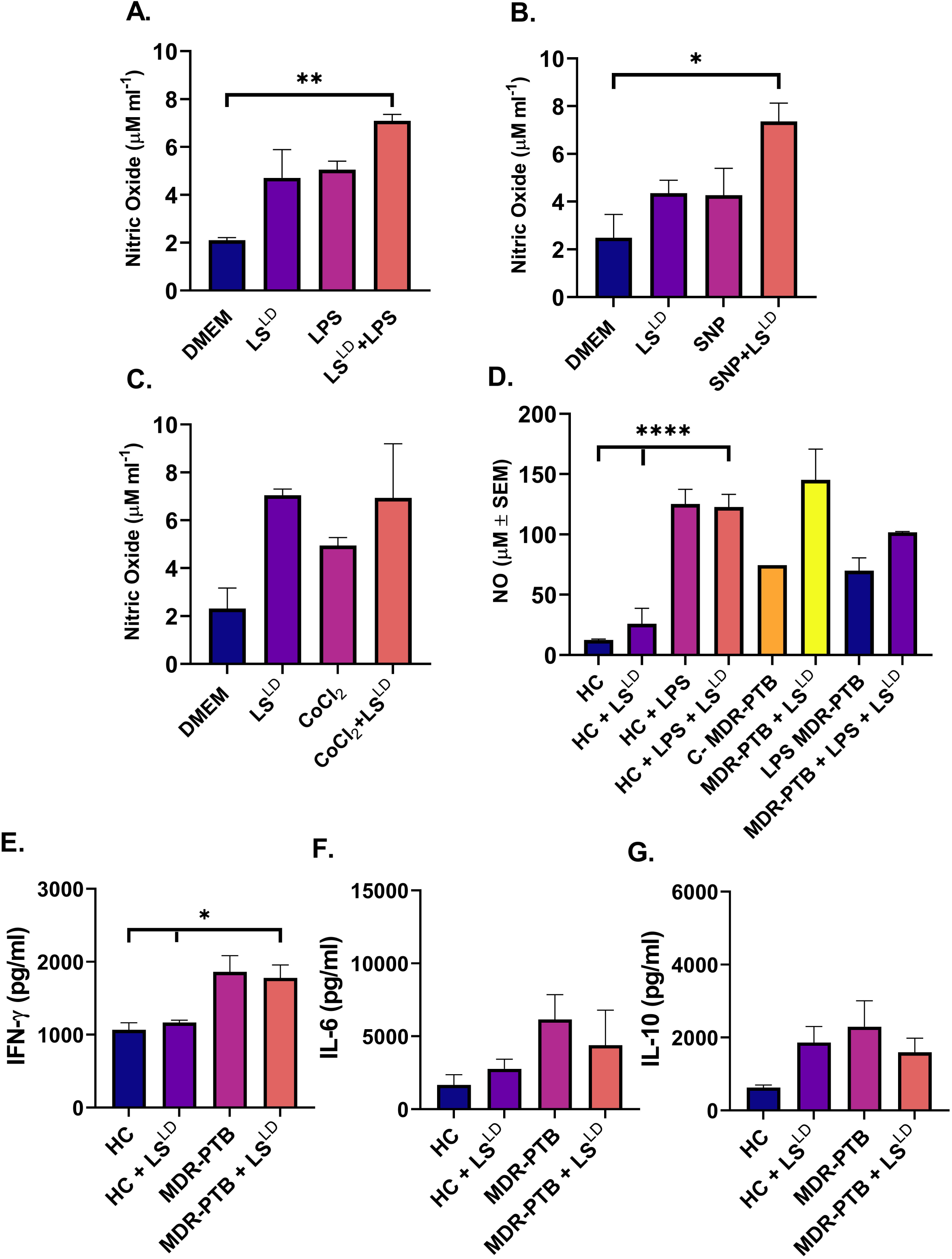
LS modulates NO and cytokine production in *in-vitro* model and patient samples. **(A-C)** NO production in RAW264.7 culture across treatment conditions, with LS alone and in combination with **(A)** LPS, **(B)** SNP, **(C)** CoCl_2_. **(D-G)** CD14^+^ monocytes were isolated from peripheral blood of healthy controls and MDR pulmonary tuberculosis (MDR-PTB) patients, differentiated into macrophages, and stimulated with indicated treatments. **(D)** NO production in CD14^+^ monocytes culture from healthy controls and MDR-PTB patients. **(E-G)** Quantitative analyses of cytokine release in CD14^+^ monocytes supernatants from healthy controls and MDR-PTB patients for IFN-γ, IL-6, and IL-10 respectively. Data are presented as mean ± SD of NO (µM) or cytokine concentrations (pg/mL) from CD14⁺ monocytes isolated from five healthy controls and five MDR-PTB patients.

Next, we estimated the NO levels in primary human CD14+ monocytes obtained from healthy controls (HC) and MDR-PTB patients. LS markedly increased both basal and LPS induced NO production in monocytes obtained from HC and MDR-PTB (**Figure 2D)**. Additionally, elevated NO production was associated with elevated IFN-γ secretion (**Figure 2E)**. Simultaneously, we observed decline in IL-6 and IL-10 levels in CD14 + monocytes from MDR-PTB (**Figure 2F-G)**. This cytokine profile aligns with increased macrophage activation and pro-inflammatory immune condition conducive to mycobacterial elimination. Both LS and palmitoyl-CoA utilized as substrates in de novo sphingolipid biosynthesis pathway. So, next we investigated the involvement of sphingolipid metabolic pathways during LS host mediated adjunctive therapy. LS mono-treatment upregulated the expression of key enzymes involved in sphingolipid pathway e.g., SPHK1, and CERS1 (**Figure 3A–E, Supplementary Table 3)**. LS and PCoA dual treatments also substantially upregulated the expression of sphingomyelin– ceramide biosynthesis, including SPTLC2 and CERS1 in HC and MDR-PTB patients (**Figure 4A–E, Supplementary Table 3)**. The activation of these pathways provides a mechanistic foundation for the increased antimycobacterial efficacy observed after LS supplementation. In conclusion, ceramide and associated sphingolipid intermediates regulate NO generation, inflammatory signalling, phagolysosomal development and various host defence mechanisms crucial for limiting intracellular *M. tuberculosis*.

**Figure 3.**
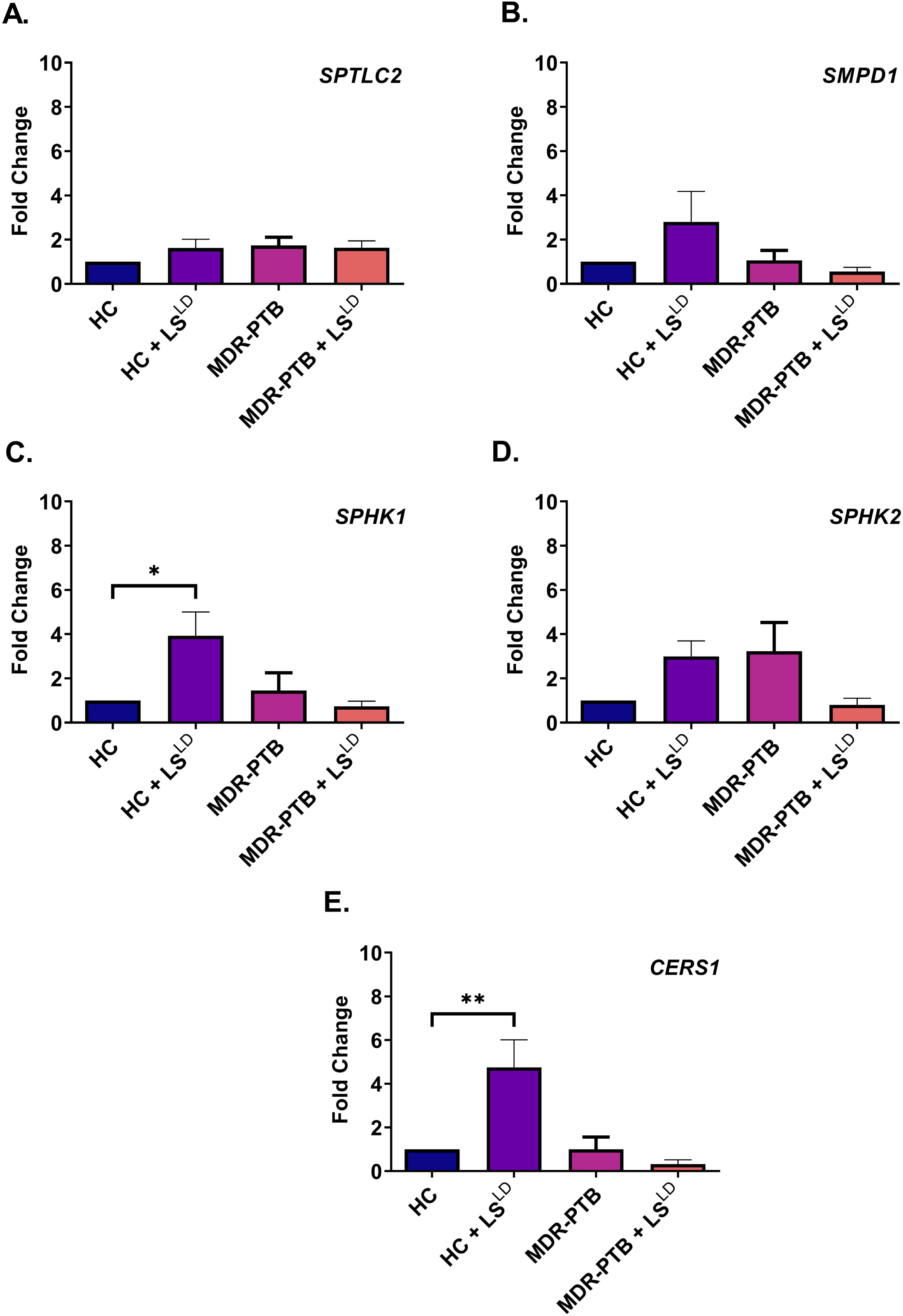
LS and PA modulate oxygen-dependent sphingolipid metabolic pathways in human CD14⁺ monocytes. **(A-E)** CD14^+^ monocytes isolated from healthy controls and MDR-PTB patients and stimulated with LS alone. qRT-PCR was performed to assess the expression of key sphingolipid pathway genes, including, SPTLC2, SMPD1, SPHK1, SPHK2, and CERS1 respectively. Fold change was calculated using the 2-ΔΔCt method and normalized to β-actin. Data represent gene expression profiles obtained from six healthy controls and six MDR-PTB patients.

**Figure 4.**
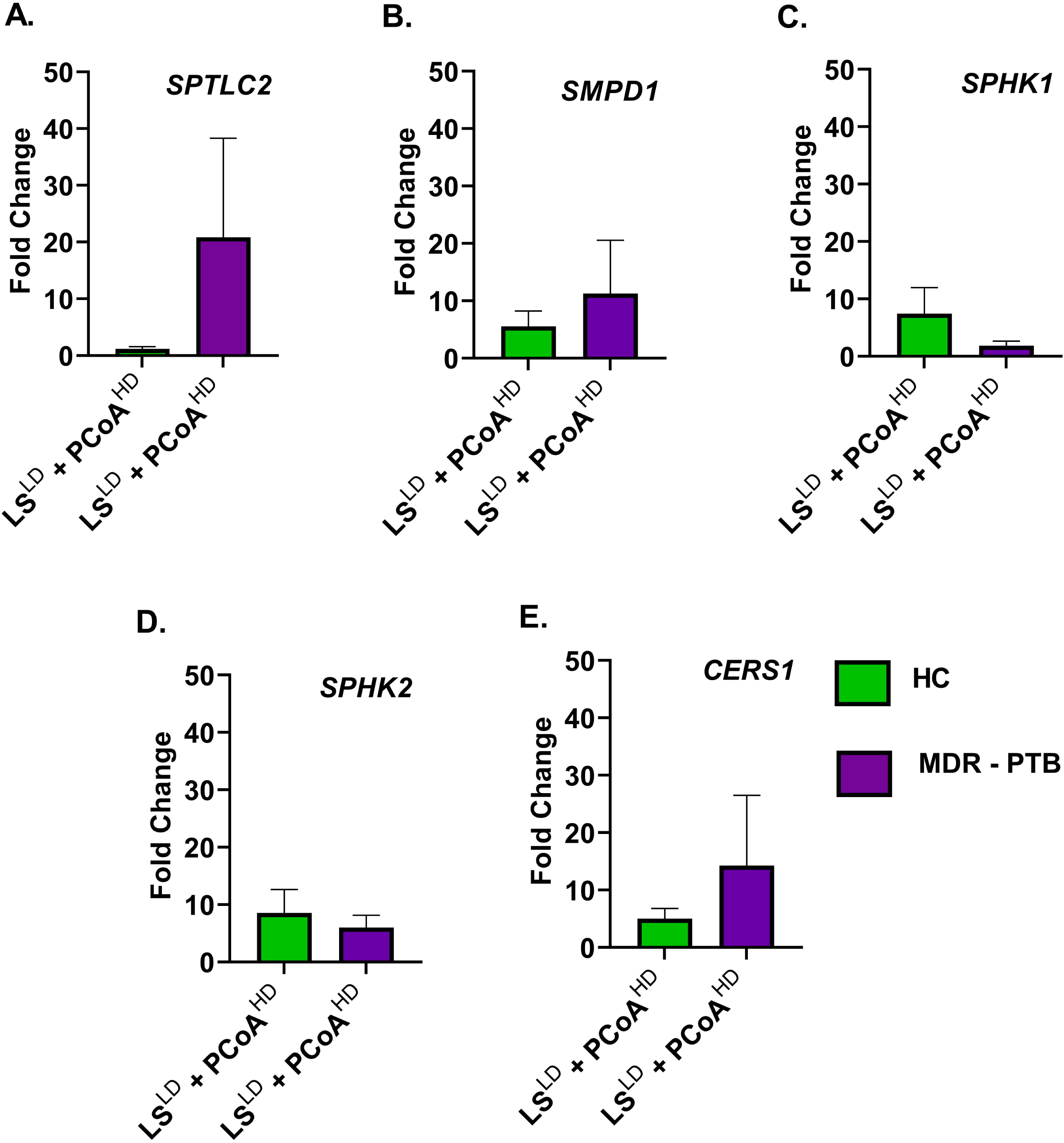
Combination of LS and PCoA modulate sphingolipid biosynthesis pathway. **(A-E)** CD14^+^ monocytes were isolated from healthy controls and MDR-PTB patients and stimulated with LS in combination with PCoA. qRT-PCR validation including genes SPTLC2 and CERS1 involved in sphingomyelin-ceramide biosynthesis. Fold change was calculated using the 2-ΔΔCt method and normalized to β-actin. Data represents gene expression profiles obtained from six healthy controls and six MDR-PTB patients.

### L-Serine and Palmitic acid Regulate Foamy Macrophage Formation

The foamy macrophage generation is the key hallmark during the tuberculosis progression. *M.tb* sends the signals to the host in order to accumulate the excess lipids, which are essential for long term survival and provide a suitable niche for the pathogen. Next we investigated the foamy macrophage formation in MDR-PTB patients and estimated the NO levels. PBMCs were obtained from both healthy individuals and MDR patients, further treated with M-CSF for macrophage maturation. Different lipid treatments (Oleic acid, naïveLDL, OxyLDL and PA) in combination with LS was given to both groups and lipid accumulation was estimated in the matured macrophages. Differentiated macrophages formed distinctive internal lipid droplets accumulation in response to additional lipid-inducing stimuli, indicating the formation of foamy macrophages. Although, lipid accumulation was observed in both healthy and MDR groups, however, LS treatment reduces the lipid accumulation **(Supplementary Figure 3)**. We also quantified the foamy macrophage formation and found that LS treatment reduced the foamy phenotype in combination with different lipids in healthy controls. Similarly, foamy macrophage formation was diminished by administration of LS in MDR patients (**Figure 5C)**. Increased foam cell production in MDR patients’ macrophages indicates altered immune activation and lipid metabolism, which reflects the pathogen-driven modification of host lipid pathways during infection. Further we measured the NO levels during different lipid treatments in healthy and MDR patients, since NO levels correlate with the functional plasticity of macrophages.

**Figure 5.**
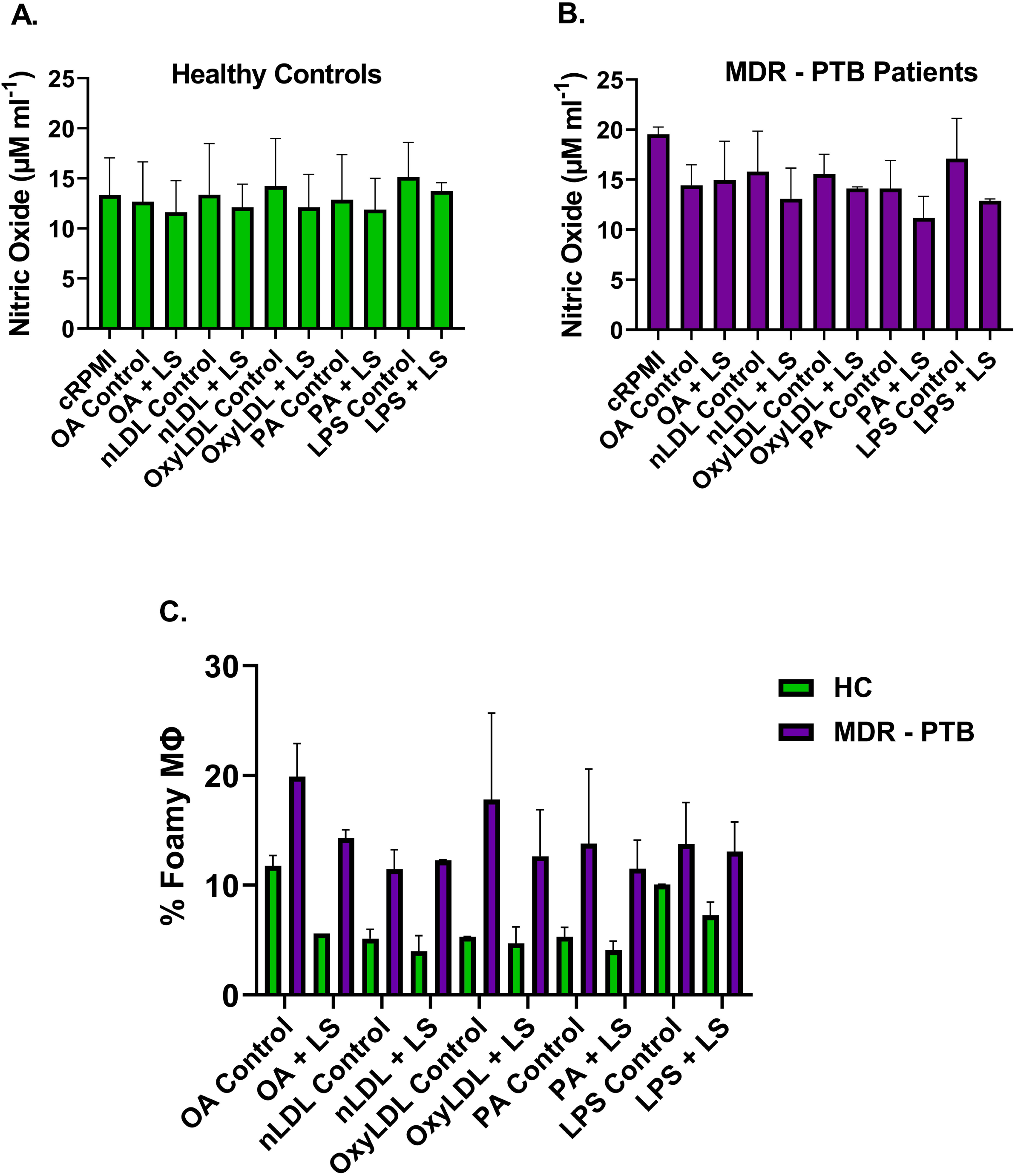
Reduction in lipid-induced foamy macrophage formation upon LS treatment in CD14^+^ monocytes. **(A-B)** NO production in CD14^+^ monocytes from **(A)** healthy control and **(B)** MDR-PTB patients stimulated with various lipid conditions with and without LS. **(C)** Percentage of foamy macrophages in healthy control and MDR-PTB patients following lipid stimulation in the presence and absence of LS.

Foamy macrophages are believed to be Th2 in nature, these macrophages normally produce less NO in comparison to normal macrophages, when stimulated in a TLR dependent manner. The observed variation in NO production between the MDR-TB patients and healthy groups show differences in immune response states and macrophage activation. While treatment with L-Serine regulated NO levels, indicating alterations in macrophage activation, lipid stimulation dramatically changed the generation of NO in macrophages (**Figure 5A-B)**. In macrophages from TB patients, decreased or altered NO levels indicate impaired antimicrobial activity, which reflects the pathogen’s capacity to alter host defence mechanisms and affect the course of the disease. These macrophages contribute to the pathogenic response in TB patients and carry scavenging phenotypes.

### L-Serine and Palmitic Acid Promote Clearance of Mycobacterial Burden in Animal Models

Based on the robust anti-tubercular efficacy of LS and palmitoyl CoA observed in macrophage infection models. We hypothesized that both metabolites would similarly reduce mycobacterial burden *in-vivo*. PA was examined in place of palmitoyl-CoA because PA has superior stability, while on other hand palmitoyl-CoA undergoes rapid degradation in tissues. We evaluated the two different treatment groups; a low-dose (LD) regimen consisting of LS (600 mg/kg) + PA (150 mg/kg), and a high-dose (HD) regimen comprising: LS (1200 mg/kg) + PA (300 mg/kg). Animals were infected and treated according to the experimental protocol shown in **Figure 6A**. Mycobacterial burden was enumerated by colony-forming unit (CFU) in the lungs (**Figure 6B)** and spleens (**Figure 6C)** eight weeks post-infection. Infected untreated animals (H_37_Rv) exhibited substantially higher bacterial loads in both organs. Meanwhile, the lungs harboring the greatest burden, which is consistent with pulmonary tropism of *M.tb*. Standard anti-tubercular therapy H+R significantly reduced CFU counts in both the lungs and spleens compared with untreated infected controls.

**Figure 6.**
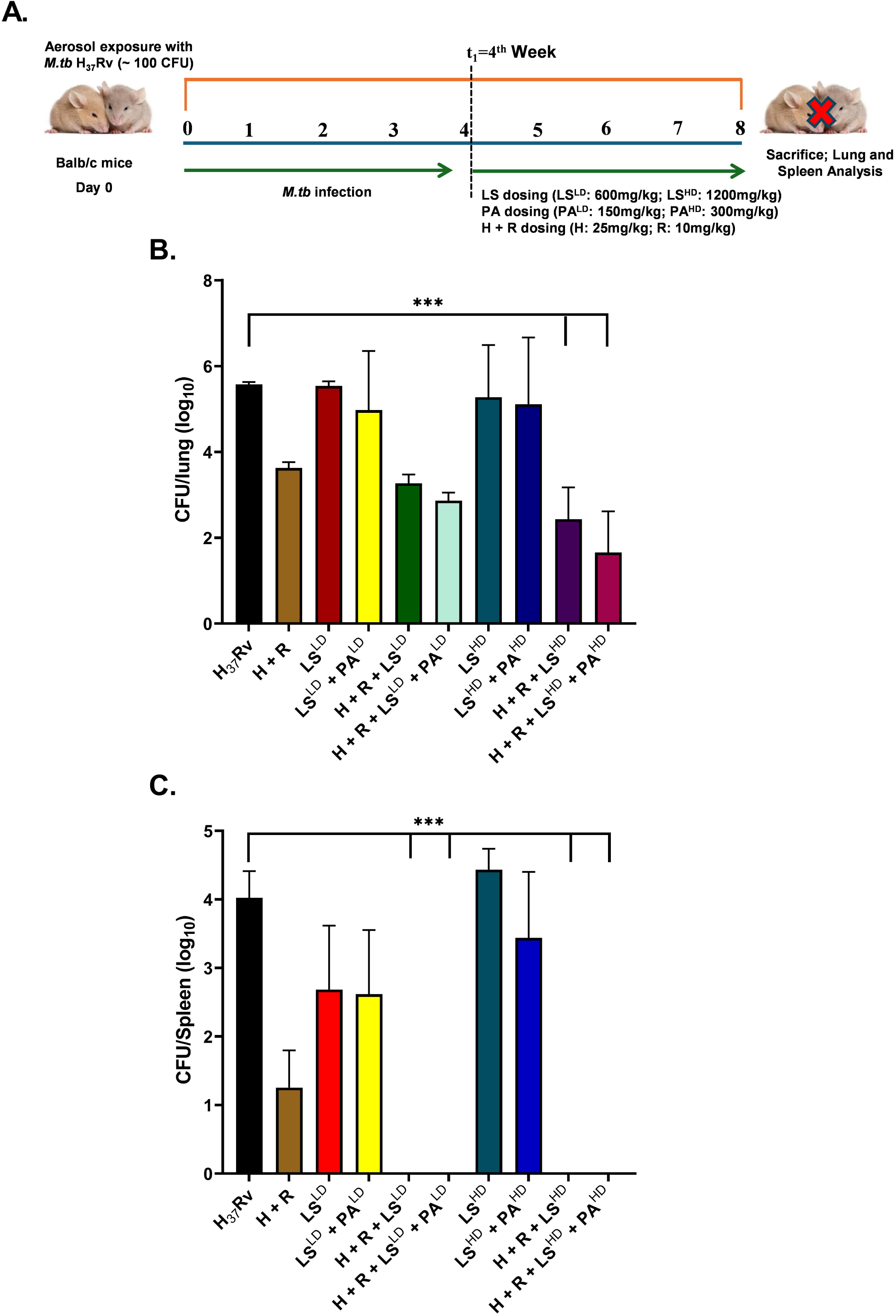
LS and PA treatment reduces pulmonary and splenic mycobacterial load in *M.tb* H37Rv infected mice. **(A)** Schematic representation of Balb/c mice were infected *with M.tb* H_37_Rv through aerosol. From the 4^th^ week onwards, LS, PA, and anti-tubercular drugs alone or combination treatment were given until 8^th^ week. Mice were euthanized after 8^th^ week post-infection to analyze the mycobacterial burden. Eight weeks post infection, lung and spleen homogenates were plated, and the mycobacterial load was measured using CFU in the **(B)** lungs and **(C)** spleens. Bacterial burdens in both organs were much higher in infected untreated mice (H_37_Rv). A two-tailed unpaired Student’s t-test was used for performing pairwise statistical comparisons *** p < 0.01 show statistical significance in relation to the infection control or between any of the specified groups.

LS monotherapy decreased mycobacterial burden in a dose-dependent manner in both organs, with the high-dose regimen producing a significant reduction in bacterial load. In contrast, PA alone exerted only a modest effect on bacterial clearance. However, co-administration of LS and PA significantly enhanced mycobacterial clearance compared with either treatment alone. Moreover, combining LS with H+R resulted in a significantly greater reduction in bacterial burden in both the lungs and spleens than H+R alone. Notably, adjunctive treatment with LS and PA together with H+R produced a further significant reduction in pulmonary CFUs compared with H+R monotreatment (p < 0.05). In the spleen, no culturable bacteria were detected in mice receiving the combination treatments, indicating complete suppression of detectable systemic dissemination.

## L-Serine and Palmitic Acid Restore Lung and Splenic Architecture in *M. tuberculosis*-Infected Mice

Next we investigated the impact of LS and PA on lung and spleen architecture during the course of *M.tb* infection. We performed histopathological analysis of lung and spleen in different treatment groups. A simultaneous dose-dependent therapeutic response was observed in both the lungs and spleen after the treatment with LS and PA, administered in combination with isoniazid and rifampicin. *M.tb* infected mice exhibited severe organ pathology, which was characterized by significant granulomatous inflammation, lung fibrosis, necrosis and extensive pulmonary consolidation **(Supplementary Figure 4; Supplementary Table 2)**. We also observed pronounced splenomegaly and substantial disruption of white and red pulp architecture that was indicative of progressive local infection and systemic immune dysfunction **(Supplementary Figure 5)**. The combination of isoniazid and rifampicin treatment improved the amelioration of pathological alteration in both organs. The histopathology showed decreased pulmonary lesions and splenic enlargement; nonetheless residual inflammation and structural abnormalities remain **(Supplementary Figures 4C and 5C; Supplementary Table 2)**. LS treatment offered modest, dose-dependent protection, leading to substantial improvement in pulmonary and splenic histopathology. However, inflammatory infiltrates, fibrosis, and architectural distortion were still apparent, especially at the lower LS dose. The combinatorial treatment of PA and LS further improved the tissue recovery, suggesting an additive impact on organ repair and restoration of cellular homeostasis.

The triple combination therapy (LS + isoniazid + rifampicin) resulted in significant improvements in both pulmonary and splenic histopathology. We observed the high dose of LS (1200 mg/kg) greatly potentiated the therapeutic efficacy of isoniazid and rifampicin in significant manner. This was evident by a marked decrease in granulomatous inflammation, maintenance of alveolar architecture, reduction of splenomegaly, and restoration of normal splenic white and red pulp organization **(Supplementary Figures 4E and 5I; Supplementary Table 2)**. The highest therapeutic effect was attained with the combination of high-dose LS, PA, isoniazid and rifampicin, leading to nearly complete histological recovery in both organs **(Supplementary Figures 4G and 5K; Supplementary Table 2)**. The lung samples showed maintained alveolar architecture without any observable granulomatous lesions, necrosis, fibrosis, or inflammatory infiltrates. Similarly, the spleens exhibited unaltered white and red pulp architecture without evidence of inflammation or fibrosis, closely akin to those of naïve control animals. These results demonstrate that LS and PA significantly improve isoniazid and rifampicin driven resolution of tuberculosis-associated tissue pathology, facilitating structural restoration in both pulmonary and systemic immune compartments.

## L-Serine and Palmitic Acid Maintain an Immunogenic Pulmonary Microenvironment in Infected Animals

Immune homeostasis is the direct reflection of disease severity and treatment response. We quantified two prominent pro- and anti-inflammatory (TNF-α and IL-10) cytokines to gauge the treatment response and disease severity in lung microenvironment. We measured the cytokine levels during different doses of LS and PA in combination with isoniazid and rifampicin. We found that pulmonary cytokine levels were influenced by LS and PA as host immune response in dose dependent manner. (**Figure 7A–D)**. Low-doses of LS and PA significantly enhanced TNF-α production relative to isoniazid and rifampicin combination, while LS monotreatment decreased the TNF-α production (**Figure 7A)**. In contrast, the elevated doses of LS and PA, especially when used in combination with isoniazid and rifampicin significantly decreased the TNF-α production (**Figure 7B)**.

**Figure 7.**
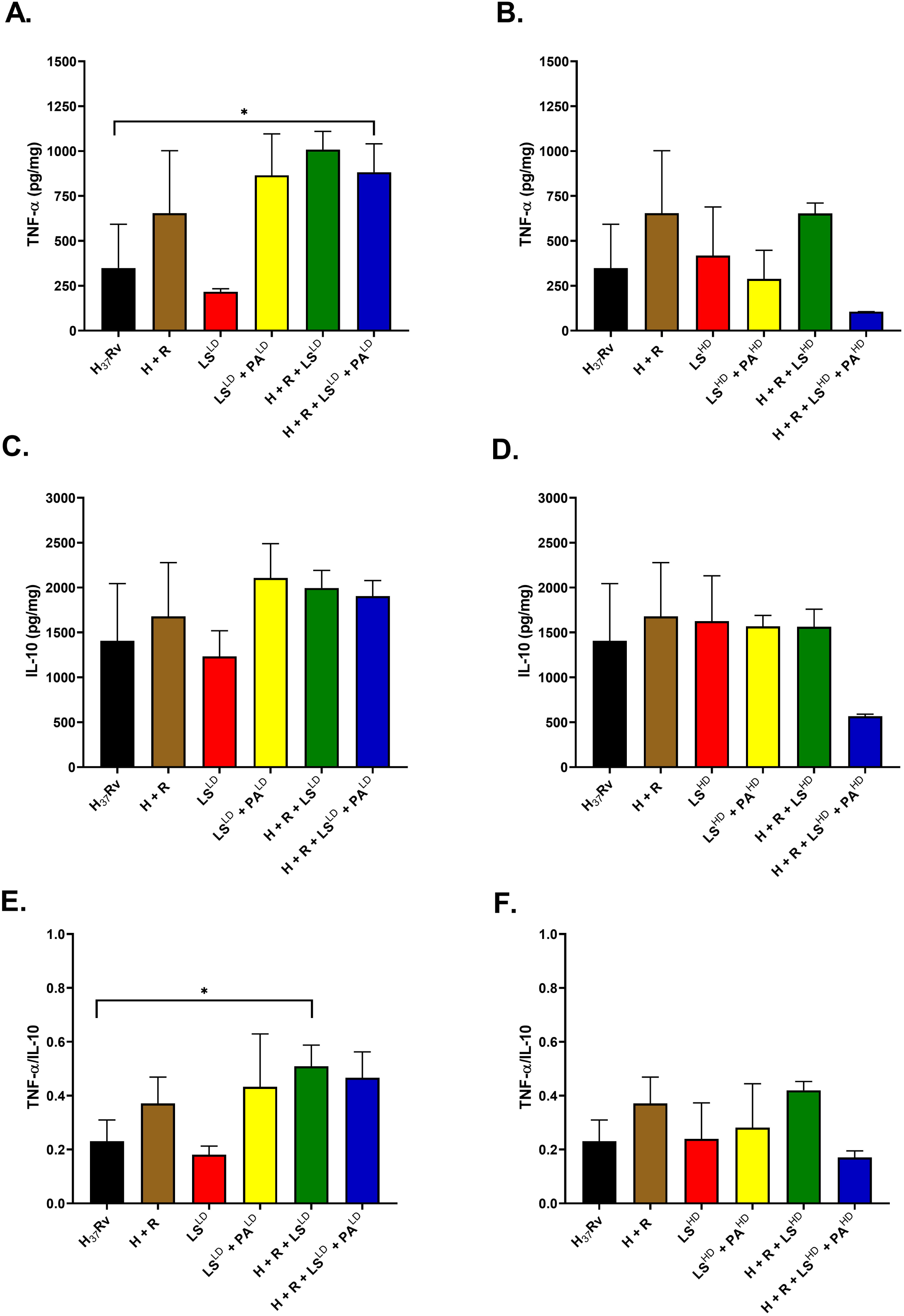
LS and PA regulate pulmonary cytokine responses in mice infected with *M.tb*. Lung homogenates were estimated for concentrations of TNF-α and IL-10 via ELISA (pg/mg protein). Pro-Inflammatory cytokine TNF-alpha concentration was measured in **(A)** Low dose LS and **(B)** High dose LS treated mice. The anti-inflammatory cytokine IL-10 concentration was measured in **(C)** Low dose LS and **(D)** High dose LS treated mice. **(E-F)** To ascertain the immunological balance, we evaluated the TNF-α/IL-10 ratio. The TNF-α/IL-10 ratio was markedly elevated by LS^LD^ and PA^LD^ combined with H and R, indicating a shift to a more immunogenic but controlled pulmonary immunological milieu. The ratio was much higher with the LS^HD^ and PA^HD^, although the trend was comparable to the low dose (* p < 0.05). Data are presented as mean ± SD. Statistical significance was determined using one-way ANOVA.

Further, we found a similar dose-dependent pattern for anti-inflammatory cytokine IL-10. Low-doses of LS and PA substantially elevated the IL-10 production, surpassing the levels attained with isoniazid and rifampicin or metabolites individually. In contrast, we observed the reduced IL-10 expression during high doses of LS and PA particularly in combination with isoniazid and rifampicin, decreased IL-10 expression, although addition of LS partially maintained the IL-10 levels (**Figure 7C-D)**. We measured the pro- and anti-inflammatory (TNF-α/IL-10) ratio to determine the immune equilibrium. Low-doses of LS and PA in combination with isoniazid and rifampicin significantly increased the TNF-α/IL-10 ratio, suggesting a transition to more immunogenic but regulated pulmonary immune environment. Although, the ratio was substantially increased during high doses of LS and PA, but the trend was like low dose (**Figure 7E-F)**.

Together, these results indicate that LS and PA adjust pulmonary cytokine response in a dose dependent fashion. The TNF-α/IL-10 ratio preserves a balanced inflammatory setting that likely supports efficient microbial immunity, while reducing excessive immunopathology. These findings endorse the promise of LS and PA as host mediated immunomodulatory adjunctive therapy for the tuberculosis treatment.

## Discussion

L-Serine is metabolically significant amino acid that plays physiological and neurological functions. Recent findings suggest that L-Serine influences multiple functions, such as immune cell activity, redox balance, and inflammatory signalling. Serine metabolism alterations have linked to the development of several diseases like asthma, pulmonary hypertension, respiratory infections, and pulmonary fibrosis. Moreover, experimental investigations suggest that the adjustments of L-Serine levels promote tissue healing, boost antioxidant ability and might reduce the inflammation. However, despite the promising preclinical results, additional mechanistic, extensive investigations and clinical trials are required to determine the safety, effectiveness and translational clinical applications of L-Serine [32]. LS works as precursor for glycine, cysteine, sphingolipids, phosphatidylserine, nucleotides. LS can modulate the neurotransmission mediated by N-methyl-D-aspartate (NMDA) receptors and influence synaptic plasticity. Disruption of LS metabolism causes the development of various neurological diseases like Alzheimer’s disease, Parkinson’s disease, and amyotrophic lateral sclerosis. Several preclinical studies suggest that LS supplementation boosts the neuronal bioenergetics and increases the myelin and membrane phospholipid production [33].

The serine metabolic pathway is emerging as promising target for developing the new antitubercular treatments. Since, LS is essential for *M.tb* growth, survival and virulence and works as precursor for protein synthesis, phospholipid generation, and one-carbon metabolism. In *M.tb*, the de novo synthesis of serine is driven by the sequential activities of phosphoglycerate dehydrogenase (SerA), phosphoserine aminotransferase (SerC), and phosphoserine phosphatase (SerB2). Among these SerB2 gained significant interest because it directly involves in the synthesis of L-Serine in *M.tb* [34]

Most emerging concept is growing that gut microbiota plays an active role in serine metabolism and can affect the microbial composition and host physiological response. LS level alterations have been associated with the gut disorders, e.g., inflammatory bowel disease, metabolic issues, and other inflammation-related conditions. These conditions are associated with compromised epithelial barrier function, oxidative stress, immune system dysregulation, and microbial imbalances. In contrast, supplementation of serine metabolism might boost intestinal resilience by improving epithelial repair, modulating inflammatory signalling pathways, and sustaining microbial balance. These results suggest that LS is crucial metabolic connector between dietary influences, gut microbiota function, and the host’s intestinal health [35]. Conversely, palmitic acid is an integral part of cellular lipid metabolism, and energy production. PA functions as an essential substrate for protein palmitoylation, phospholipid synthesis, and lipid-mediated signalling. High amounts of PA consumption have been linked with dyslipidemia, insulin resistance, non-alcoholic fatty liver disease (NAFLD), type 2 diabetes (T2D), and cardiovascular diseases (CVD) [36, 37].

Serine metabolism is also essential in regulating immune function and modulating the host microenvironment during infection. So far, the impact of serine on antimycobacterial immunity remains unexplored. A metabolic shift in the serine synthesis profoundly affects the *M.tb* survival inside the host by modulating the macrophage activation and antimicrobial response. Recently, the role of serine metabolism in *M.tb*-infected macrophages has been investigated. The study reveals that *M.tb* infection induces enzymes associated with the serine synthesis pathway (SSP) and serine transporters. Moreover, inhibition of the key SSP enzyme or restriction of exogenous serine boosts antimycobacterial immunity in both *in-vitro* and *in-vivo*. Further, experiments revealed that depletion of serine reduces reactive oxygen species (ROS) levels by diminishing the levels of reduced nicotinamide adenine dinucleotide. This ROS reduction destabilizes hypoxia-inducible factor 1-alpha (HIF-α), impairing the glucose uptake and adenosine triphosphate (ATP) generation. Consequently, reduced ATP production upregulates the adenosine monophosphate-activated protein kinase, which further inhibits mammalian target of rapamycin (mTOR) and induces autophagy thereby, exerting an antimycobacterial effect. These investigations reveal the serine’s role as a crucial immune metabolite during *M.tb* infection and propose that manipulating serine metabolism holds therapeutic promise against mycobacterial infections [38]. In another study, an untargeted metabolomics analysis showed that the serine catabolic pathway was inhibited in drug-resistant *Streptococcus suis*. LS supplementation restored the fungicidal effect of macrolides on *S. suis in-vivo* and *in-vitro* by enhancing the serine metabolic pathway. Further studies showed that L-Serine, stimulated by its serine catabolic pathway, inhibited intracellular H_2_S production, and reduced Fe-S cluster production. It also attenuated the production of glutathione, an important marker of the intracellular oxidation-reduction reaction. All these phenomena eventually contribute to an increase in the level of reactive oxygen species, which leads to intracellular DNA damage and bacterial death. This study provides a potential new approach for treating diseases caused by drug-resistant *S. suis*. [39].

In this context, our investigations recognize LS combined with isoniazid and rifampicin as an emerging HDT. This adjunctive therapy improves antimycobacterial efficacy across *in-vitro*, intracellular, and *in-vivo* models. In axenic culture experiments, LS exhibited the limited antimycobacterial efficacy, requiring minimal inhibitory concentrations in the millimolar range for the drug-susceptible *M.tb* and significantly elevated concentrations for MDR strains. These results indicate that LS is probably not clinically effective as a standalone antimicrobial agent, because of its consistent rapid systemic utilization and metabolic clearance. Nevertheless, checkerboard evaluations revealed that LS significantly enhanced the efficacy of rifampicin and moxifloxacin, even against the MDR strain, FICI values ≤0.5. Similarly, antibiotic-sensitizing effects of LS also observed in other bacterial pathogens, such as *Mycobacterium tuberculosis*, *Streptococcus suis* and *Acinetobacter baumannii*, LS decreases the effective concentrations of macrolides or directly displays the inhibitory effect and re-establishes the bactericidal activity [38–40]. Together, these observations suggest that LS mainly acts as a metabolic enhancer, that improves antibiotic effectiveness through either modulating the bacterial physiology or host cellular responses. This is recently evident that LS can exert direct bactericidal activity.

The host-mediated effects of LS were particularly evident in macrophage infection models. LS reduced intracellular mycobacterial burden in both concentration- and time-dependent manners in murine RAW264.7 macrophages, human THP-1-derived macrophages, and primary CD14⁺ monocyte-derived macrophages. These effects were consistently and significantly amplified by co-administration of PA, highlighting the significance of coordinated substrate availability for *de-novo* sphingolipid biosynthesis. Mechanistically, LS enhanced NO production and induced a cytokine profile marked by elevated levels of IFN-γ and reduced levels of IL-10 and IL-6. These immunomodulatory perturbations reflect the macrophage activation state conducive to mycobacterial control. Our results aligned with earlier investigations demonstrating the sphingolipids regulate phagolysosomal maturation, autophagy, and inflammatory signalling. Meanwhile, *M.tb* actively disrupts the host sphingolipid metabolism to promote the intracellular survival [41, 42].

Our results are supported by increasing evidence that host lipid metabolism is crucial in TB pathogenesis. Clinical studies have indicated lower sphingolipid levels in individuals with active TB, especially in those with severe illness, malnutrition and conditions associated with impaired immune function. Through replenishing the necessary substrates required for sphingolipid biosynthesis, LS and PA seem to re-establish immunometabolic homeostasis, thereby, improving the antimycobacterial defence while mitigating the negative effects linked to direct sphingosine-1-phosphate administration, such as vascular leakage and immune dysregulation. Interestingly, the balanced TNF-α/IL-10 responses observed after LS-based combination therapy indicate a dual boost of antimicrobial defence and reduction of excessive inflammation, fulfilling a central objective of HDT strategies [43].

The physiological significance of our results was further validated in a murine aerosol infection model. Although, LS monotherapy produced moderate, dose-dependent reductions in pulmonary and splenic bacterial loads, when we administered LS as adjunct with isoniazid and rifampicin significantly enhanced bacterial clearance. The highest therapeutic efficacy was attained through the combined administration of LS, PA, isoniazid and rifampicin. We observed a significant reduction in pulmonary colony-forming units (CFUs) and total eradication of culturable bacteria from the spleen. Histopathological evaluations confirmed these microbiological results, demonstrating almost total resolution of granulomatous inflammation, necrosis, fibrosis, and related tissue damage, especially at higher LS doses. The application of PA instead of palmitoyl-CoA *in-vivo* is based on practical aspects of stability and bioavailability. We believe PA endogenously converts into palmitoyl-CoA through established metabolic pathways, consistent with earlier dietary supplementation research [17, 18, 22, 23].

LS has multiple characteristics from a translational potential, making it a compelling and attractive adjunctive HDT candidate. Being a naturally occurring amino acid with an established safety profile and Generally Recognized As Safe (GRAS) status, LS is highly appropriate and ideal candidate for oral administration and incorporation into existing TB treatment regimens [44]. Through enhancing the effectiveness of antibiotics, LS-based adjunctive therapy could decrease antimicrobial dosages, can shorten the treatment duration, and improve the therapeutic outcomes in MDR -TB. Moreover, the consistent activity observed in both murine and human macrophage models underscore its translational potential. However, several important limitations should be acknowledged during translational implementations. The MDR analyses were performed on limited sets of clinical isolates and thus, we need further verification in broader collections encompassing genetically diverse MDR and XDR strains. Enhanced sphingolipid metabolism and NO production strongly correlated with improved antimicrobial outcomes. Moreover, the further investigations are required to provide cocnclusive mechanistic evidence targeting genetic or pharmacological disruption of these pathways. Additionally, the dosing schedules used in mouse studies need comprehensive pharmacokinetic and pharmacodynamic optimization prior clinical application, especially in malnourished patients with TB.

In summary, our results highlight LS, particularly in combination with PA, acts as a potential metabolic supplement for host-directed treatment of tuberculosis. Moreover, instead of functioning as a direct antimicrobial agent, LS boosts the effectiveness of primary anti-tubercular medications and encourages immunometabolic reprogramming. This two-fold mechanism is evident through enhanced bacterial elimination, re-establishment of balanced inflammatory responses, and maintenance of tissue architecture in preclinical TB models. Importantly, our research demonstrates that metabolic supplementation with LS and PA can reprogram host cellular pathways, such as antimicrobial defence, creating a mechanistic connection between nutrient accessibility, sphingolipid metabolism, and the management of intracellular pathogens. Our findings offer both mechanistic understanding and a compelling translational justification for developing nutrient-focused host-directed therapies to improve treatment outcomes in drug-susceptible and multidrug-resistant tuberculosis.

## Acknowledgment

We thank all the study participants and the staff of the Staff of ICMR-National Jalma Institute of Leprosy & Other Mycobacterial Diseases, Agra for helping us in the animal challenge experiments. We also express gratitude the study participants and the staff members from Department of Pulmonary Medicine, All India Institute of Medical Sciences, New Delhi. We acknowledge Dr. Showket Hussain, ICMR-National Institute of Cancer Prevention and Research, Noida, Uttar Pradesh for providing help in real time experiments. We thank Ms. Vandana Mehra from Amity Centre for Translational Research, Amity University, for providing critical comments on the manuscript.

## Ethical Approval

The study received approval from the Institutional Ethics Committees (IEC) of Amity University, Noida (AUUP/IEC/May 2023/7 IEC). The study protocol was approved for animal experiments from the Institutional Biosafety Committee, Institutional Animal Ethics Committee, ICMR- National Jalma Institute of Leprosy & Other Mycobacterial Diseases, Agra (NJIL&OMD/61AEC /2022-05), and Institutional Human Ethics Committee, All India Institute of Medical Sciences, New Delhi (AIIMS/REV2023-44). Written informed consent was obtained from all healthy controls and patients with tuberculosis prior to sample collection. Studies in this work abide by the Declaration of Helsinki principles.

## Sequence Information

Not applicable.

## Data Availability

The data supporting the findings of this study are available from the corresponding author upon reasonable request. Data generated from CD14⁺ monocytes isolated from healthy controls and patients with pulmonary tuberculosis (PTB) are not publicly available because they contain sensitive human participant information.

## Author Contributions

H.P. and A.K.S. conceived and designed the study and planned the experiments. N.S., R.S., A.K. performed the *in-vitro* and *in-vivo* experiments. V.H. supervised patient recruitment and the prospective clinical study involving CD14⁺ monocytes from healthy controls and PTB patients and contributed to manuscript editing. A.A. performed the histopathological analyses. H.P., A.K.S., A.K. and L.K.S. conducted data analysis, visualization of data, and provided methodological guidance. H.P., A.K.S. and L.K.S. prepared the original draft of the manuscript. All authors reviewed, revised, and approved the final version of the manuscript.

## References

1. Global Tuberculosis Report 2025. World Health Organization. ISBN 978–92-4- 011693-1, https://www.who.int/publications/i/item/9789240116924.

2. Rolando, M. and Buchrieser, C., 2019. A comprehensive review on the manipulation of the sphingolipid pathway by pathogenic bacteria. Frontiers in Cell and Developmental Biology, 7, p.168. doi: 10.3389/fcell.2021.647045.

3. Mohan, M. and Bhattacharya, D., 2021. Host-directed therapy: a new arsenal to come. Combinatorial Chemistry & High Throughput Screening, 24(1), pp.59–70. doi: 10.2174/1386207323999200728115857.

4. Zhang, Y., Wu, R., Sun, M., Li, X., Fang, R., Xing, J., Li, Z., Wen, Y. and Song, N., 2025. Progress of anti-tuberculosis drug targets and novel therapeutic strategies. Frontiers in Microbiology, 16, p.1637254. doi: 10.3389/fmicb.2025.1637254. Paton NI, Cousins C, Suresh C, Burhan E, Chew KL, Dalay VB, Lu Q, Kusmiati T, Balanag VM, Lee SL, Ruslami R. Treatment strategy for rifampin-susceptible tuberculosis. New England Journal of Medicine. 2023 Mar 9;388(10):873-87. doi: 10.1056/NEJMoa2212537.

5. Tian, N., Chu, H., Li, Q., Sun, H., Zhang, J., Chu, N. and Sun, Z., 2025. Host-directed therapy for tuberculosis. European Journal of Medical Research, 30(1), p.267. doi: 10.1186/s40001-025-02443-4.

6. Lee, M., Lee, S.Y. and Bae, Y.S., 2023. Functional roles of sphingolipids in immunity and their implication in disease. Experimental & Molecular Medicine, 55(6), p.1110–1130. doi: 10.1038/s12276-023-01018-9.

7. Seitz, A.P., Grassmé, H., Edwards, M.J., Pewzner-Jung, Y. and Gulbins, E., 2015. Ceramide and sphingosine in pulmonary infections. Biological Chemistry, 396(6-7), p.611–620. doi: 10.1515/hsz-2014-0285.

8. Chen, X., Ye, J., Lei, H. and Wang, C., 2022. Novel potential diagnostic serum biomarkers of metabolomics in osteoarticular tuberculosis patients: A preliminary study. Frontiers in Cellular and Infection Microbiology, 12, p.827528. doi: 10.3389/fcimb.2022.827528.

9. Nadella, V., Sharma, L., Kumar, P., Gupta, P., Gupta, U.D., Tripathi, S., Pothani, S., Qadri, S.S.Y.H. and Prakash, H., 2020. Sphingosine-1-phosphate (S-1P) promotes differentiation of naïve macrophages and enhances protective immunity against *Mycobacterium tuberculosis*. Frontiers in Immunology, 10, p.3085. doi: 10.3389/fimmu.2019.03085.

10. Sharma, L. and Prakash, H., 2017. Sphingolipids are dual specific drug targets for the management of pulmonary infections: perspective. Frontiers in Immunology, 8, p.378. doi: 10.3389/fimmu.2017.00378.

11. Gutierrez, M.G., Gonzalez, A.P., Anes, E. and Griffiths, G., 2009. Role of lipids in killing mycobacteria by macrophages: evidence for NF-κB-dependent and-independent killing induced by different lipids. Cellular Microbiology, 11(3), p.406–420. doi: 10.1111/j.1462-5822.2008.01263.x.

12. Sun, G., Wang, B., Wu, X., Cheng, J., Ye, J., Wang, C., Zhu, H. and Liu, X., 2024. How do sphingosine-1-phosphate affect immune cells to resolve inflammation? Frontiers in Immunology, 15, p.1362459. doi: 10.3389/fimmu.2024.1362459.

13. Niekamp, P., Guzman, G., Leier, H.C., Rashidfarrokhi, A., Richina, V., Pott, F., Barisch, C., Holthuis, J.C. and Tafesse, F.G., 2021. Sphingomyelin biosynthesis is essential for phagocytic signaling during *Mycobacterium tuberculosis* host cell entry. mBio, 12(1), p. e03141–20. doi: 10.1128/mBio.03141-20.

14. Singh, A.K., Gangakhedkar, R., Thakur, H.S., Raman, S.K., Patil, S.A. and Jain, V., 2023. Mycobacteriophage D29 Lysin B exhibits promising anti-mycobacterial activity against drug-resistant *Mycobacterium tuberculosis*. Microbiology Spectrum, 11(6), p.e04597–22. doi: 10.1128/spectrum.04597-22.

15. Layoun, A., Samba, M. and Santos, M.M., 2015. Isolation of murine peritoneal macrophages to carry out gene expression analysis upon Toll-like receptors stimulation. Journal of Visualized Experiments, (98), p.e52749. doi: 10.3791/52749.

16. Bailey, J.D., Shaw, A., McNeill, E., Nicol, T., Diotallevi, M., Chuaiphichai, S., Patel, J., Hale, A., Channon, K.M. and Crabtree, M.J., 2020. Isolation and culture of murine bone marrow-derived macrophages for nitric oxide and redox biology. Nitric Oxide, 100, p.17–29. doi: 10.1016/j.niox.2020.04.005.

17. Singh, A.K., Yadav, A.B., Garg, R. and Misra, A., 2014. Single nucleotide polymorphic macrophage cytokine regulation by *Mycobacterium tuberculosis* and drug treatment. Pharmacogenomics, 15(4), p.497–508. doi: 10.2217/pgs.13.240.

18. Schreiber, T., Ehlers, S., Aly, S., Hölscher, A., Hartmann, S., Lipp, M., Lowe, J.B. and Hölscher, C., 2006. Selectin ligand-independent priming and maintenance of T cell immunity during airborne tuberculosis. The Journal of Immunology, 176(2), p.1131–1140 doi: 10.4049/jimmunol.176.2.1131.

19. Tada, Y., Yano, N., Takahashi, H., Yuzawa, K., Ando, H., Kubo, Y., Nagasawa, A., Chin, K., Kawamata, Y., Sakai, R. and Ohashi, N., 2010. A 90-day feeding toxicity study of L-Serine in male and female Fischer 344 rats. Journal of Toxicologic Pathology, 23(1), p.39–47. doi: 10.1293/tox.23.39.

20. Kaneko, I., Han, L., Liu, T., Li, J., Zhao, Y., Li, C., Yi, Y., Liang, A. and Hayamizu, K., 2009. A 13-week subchronic oral toxicity study of L-Serine in rats. Food and Chemical Toxicology, 47(9), p.2356–2360. doi: 10.1016/j.fct.2009.06.030.

21. Yang, Y., Yu, Q., Li, B., Yang, Z., Zhang, S. and Yuan, F., 2023. Palmitate lipotoxicity is closely associated with the fatty acid-albumin complexes in BV-2 microglia. PloS One, 18(4), p.e0281189. doi: 10.1371/journal.pone.0281189.

22. Liu, M., Gao, B.Y., Qin, F., Wu, P.P., Shi, H.M., Luo, W., Ma, A.N., Jiang, Y.R., Xu, X.B. and Yu, L.L.L., 2012. Acute oral toxicity of 3-MCPD mono-and di-palmitic esters in Swiss mice and their cytotoxicity in NRK-52E rat kidney cells. Food and Chemical Toxicology, 50(10), p.3785–3791. doi: 10.1016/j.fct.2012.07.038.

23. Yang, Z.H., Miyahara, H. and Hatanaka, A., 2011. Chronic administration of palmitoleic acid reduces insulin resistance and hepatic lipid accumulation in KK-A^y^ Mice with genetic type 2 diabetes. Lipids in Health and Disease, 10(1), p.120. doi: 10.1186/s12944-021-01513-w.

24. Sharma, A.K., Arora, D., Singh, L.K., Gangwal, A., Sajid, A., Molle, V., Singh, Y. and Nandicoori, V.K., 2016. Serine/threonine protein phosphatase PstP of *Mycobacterium tuberculosis* is necessary for accurate cell division and survival of pathogen. Journal of Biological Chemistry, 291(46), p.24215–24230 doi: 10.1074/jbc.M116.754531.

25. Agarwal, P., Combes, T.W., Shojaee-Moradie, F., Fielding, B., Gordon, S., Mizrahi, V. and Martinez, F.O., 2020. Foam cells control *Mycobacterium tuberculosis* infection. Frontiers in Microbiology, 11, p.1394. doi: 10.3389/fmicb.2020.594142.

26. Caocci, M., Niu, M., Fox, H.S. and Burdo, T.H., 2024. HIV infection drives foam cell formation via NLRP3 inflammasome activation. International Journal of Molecular Sciences, 25(4), p.2367. doi: 10.3390/ijms25042367.

27. van der Bruggen, T., Nijenhuis, S., van Raaij, E., Verhoef, J. and Sweder van Asbeck, B., 1999. Lipopolysaccharide-induced tumor necrosis factor alpha production by human monocytes involves the raf-1/MEK1-MEK2/ERK1-ERK2 pathway. Infection and Immunity, 67(8), p.3824–3829 doi: 10.1128/IAI.67.8.3824-3829.1999.

28. Al-Rashed, F., Haddad, D., Al Madhoun, A., Sindhu, S., Jacob, T., Kochumon, S., Obeid, L.M., Al-Mulla, F., Hannun, Y.A. and Ahmad, R., 2023. ACSL1 is a key regulator of inflammatory and macrophage foaming induced by short-term palmitate exposure or acute high-fat feeding. iScience, 26(7). doi: 10.1016/j.isci.2023.107145.

29. Xu, S., Huang, Y., Xie, Y., Lan, T., Le, K., Chen, J., Chen, S., Gao, S., Xu, X., Shen, X. and Huang, H., 2010. Evaluation of foam cell formation in cultured macrophages: an improved method with Oil Red O staining and DiI-oxLDL uptake. Cytotechnology, 62(5), p.473–481 doi: 10.1007/s10616-010-9290-0.

30. Sanda, G.M., Stancu, C.S., Deleanu, M., Toma, L., Niculescu, L.S. and Sima, A.V., 2021. Aggregated LDL turn human macrophages into foam cells and induce mitochondrial dysfunction without triggering oxidative or endoplasmic reticulum stress. PLoS One, 16(1), p.e0245797. doi: 10.1371/journal.pone.0245797.

31. Du, J., Zhao, L., Kang, Q., He, Y. and Bi, Y., 2023. An optimized method for Oil Red O staining with the salicylic acid ethanol solution. Adipocyte, 12(1), p.2179334. doi: 10.1080/21623945.2023.2179334.

32. Li, P., Wu, X., Huang, Y., Qin, R., Xiong, P. and Qiu, Y., 2025. L-Serine metabolic regulation and host respiratory homeostasis. Frontiers in Cellular and Infection Microbiology, 15, p.1518659. doi: 10.3389/fcimb.2025.1518659.

33. Phone Myint, S.M.M. and Sun, L.Y., 2023. L-Serine: neurological implications and therapeutic potential. Biomedicines, 11(8), p.2117. doi: 10.3390/biomedicines11082117.

34. Haufroid, M. and Wouters, J., 2019. Targeting the serine pathway: a promising approach against tuberculosis? Pharmaceuticals, 12(2), p.66. doi:10.3390/ ph12020066.

35. Devaux, A., Boucher, D., Villéger, R. and Bonnet, M., 2026. L-Serine at the crossroads of microbiota, intestinal health, and disorders. Communications Biology, 9(1), p.632. doi: 10.1038/s42003-026-10133-y.

36. Murru, E., Manca, C., Carta, G. and Banni, S., 2022. Impact of dietary palmitic acid on lipid metabolism. Frontiers in Nutrition, 9, p.861664. doi: 10.3389/fnut.2022.861664.

37. Carta, G., Murru, E., Banni, S. and Manca, C., 2017. Palmitic acid: physiological role, metabolism and nutritional implications. Frontiers in Physiology, 8, p.902. doi: 10.3389/fphys.2017.00902.

38. Son, S.H., Choi, J.A., Kim, J., Nguyen, T.D., Lee, J., Son, D., Jo, S. and Song, C.H., 2026. Targeting serine metabolism boosts antimycobacterial immunity during *Mycobacterium tuberculosis* H_37_Rv infection. Molecules and Cells, 49(7), p.100366. doi: 10.1016/j.mocell.2026.100366.

39. Wu, T., Wang, X., Dong, Y., Xing, C., Chen, X., Li, L., Dong, C. and Li, Y., 2022. Effects of L-Serine on macrolide resistance in *Streptococcus suis*. Microbiology Spectrum, 10(4), p.e00689–22. doi: 10.1128/spectrum.00689-22.

40. Zhou, J., Feng, D., Li, X., Chen, Y., Zhang, M., Wu, W., Zhu, J., Li, H., Peng, X. and Zhang, T., 2024. L-Serine enables reducing the virulence of *Acinetobacter baumannii* and modulating the SIRT1 pathway to eliminate the pathogen. Microbiology Spectrum, 12(3), p.e03226–23. doi: 10.1128/spectrum.03226-23.

41. Young, M.M. and Wang, H.G., 2018. Sphingolipids as regulators of autophagy and endocytic trafficking. Advances in Cancer Research, 140, p. 27–60. doi: 10.1016/bs.acr.2018.04.008.

42. Mohammed, S.A., Saini, R.V., Jha, A.K., Hadda, V., Singh, A.K. and Prakash, H., 2022. Sphingolipids, mycobacteria and host: Unraveling the tug of war. Frontiers in Immunology, 13, p.1003384. doi: 10.3389/fimmu.2022.1003384.

43. Cavalcanti, Y.V.N., Brelaz, M.C.A., Neves, J.K.D.A.L., Ferraz, J.C. and Pereira, V.R.A., 2012. Role of TNF-alpha, IFN-gamma, and IL-10 in the development of pulmonary tuberculosis. *Pulmonary Medicine*, 2012(1), p.745483. doi: 10.1155/2012/745483.

44. Ye, L., Sun, Y., Jiang, Z. and Wang, G., 2021. L-Serine, an endogenous amino acid, is a potential neuroprotective agent for neurological disease and injury. Frontiers in Molecular Neuroscience, 14, p.726665. doi: 10.3389/fnmol.2021.726665.

