## Supplementary Figures and Tables for "L-Serine and Palmitoyl CoA control *Mycobacterial* infection by tweaking protective immune response: A potential host directed therapy for tuberculosis"

**Running Title**
L-Serine as host-directed adjunctive therapy

#### Suppl. Figure 1

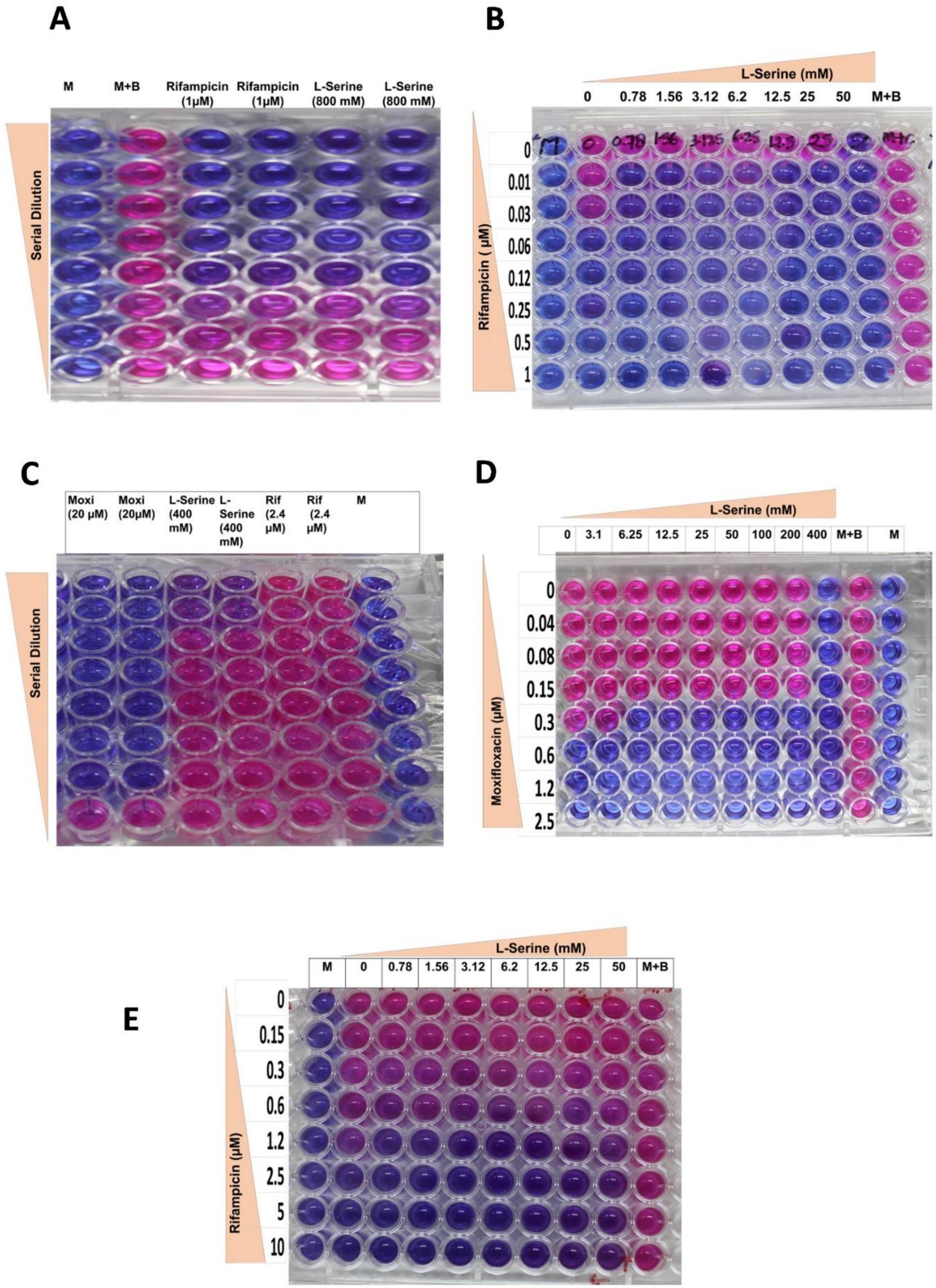

### Suppl. Figure 2

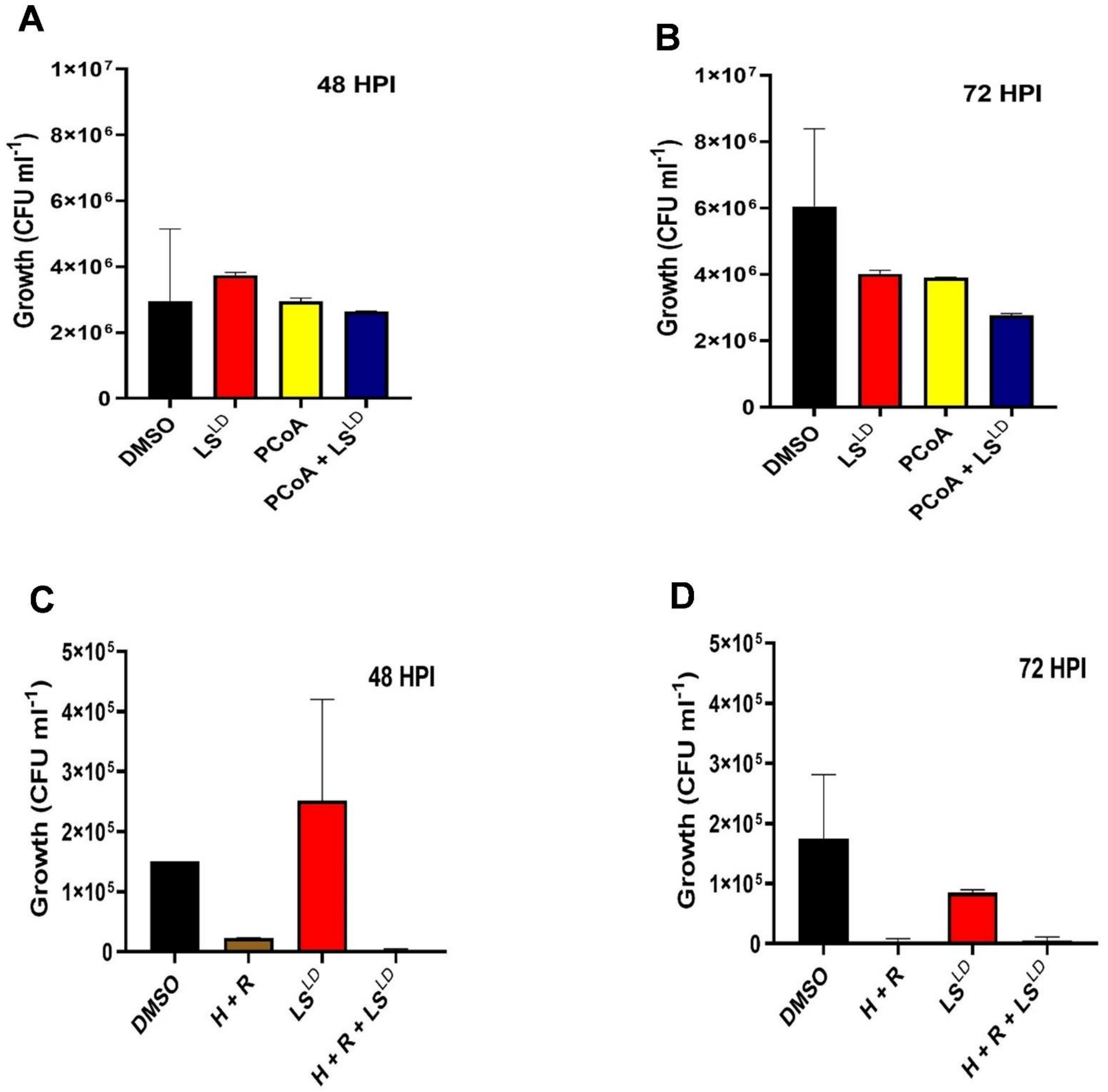

Suppl. Figure 3

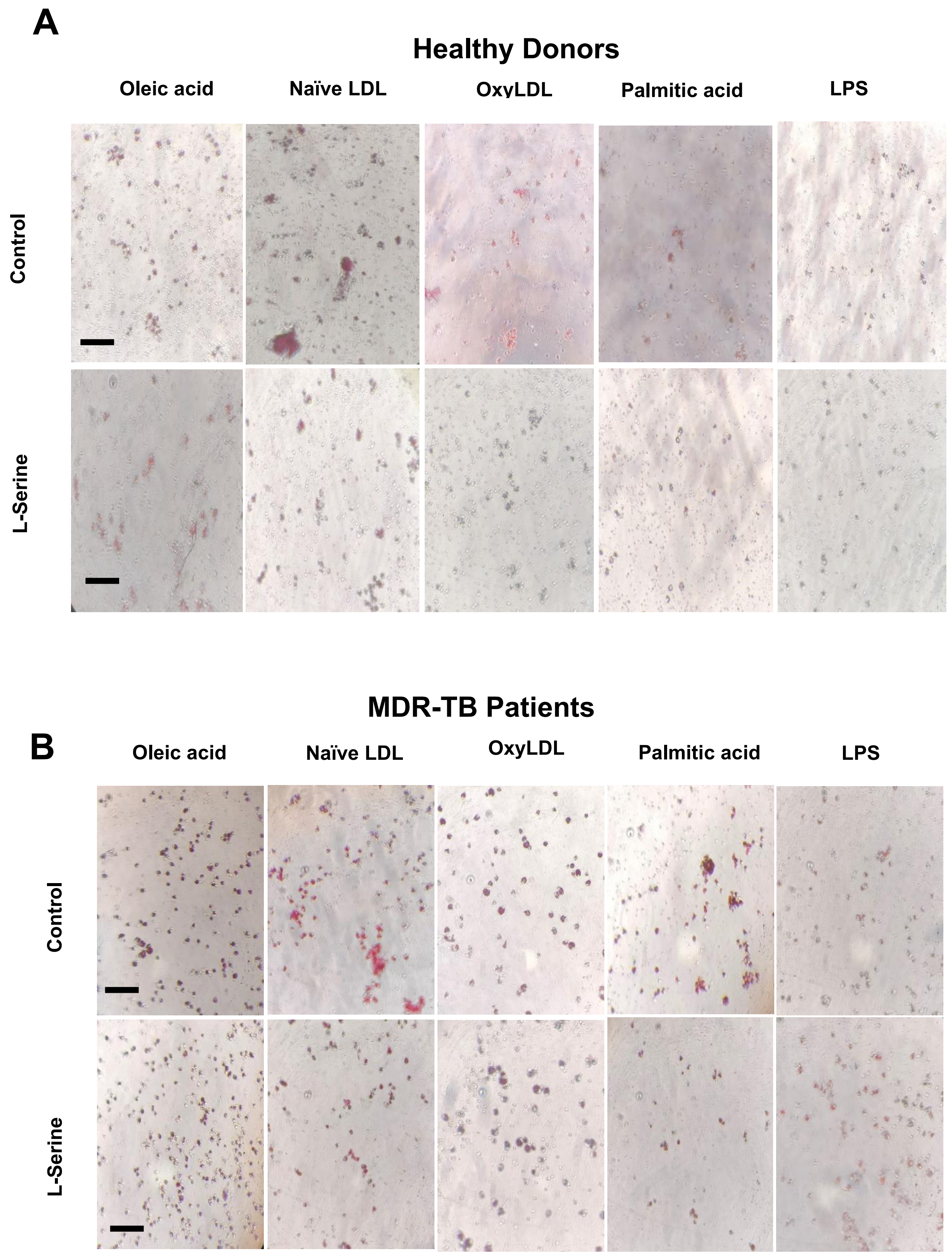

**Suppl. Figure 4**

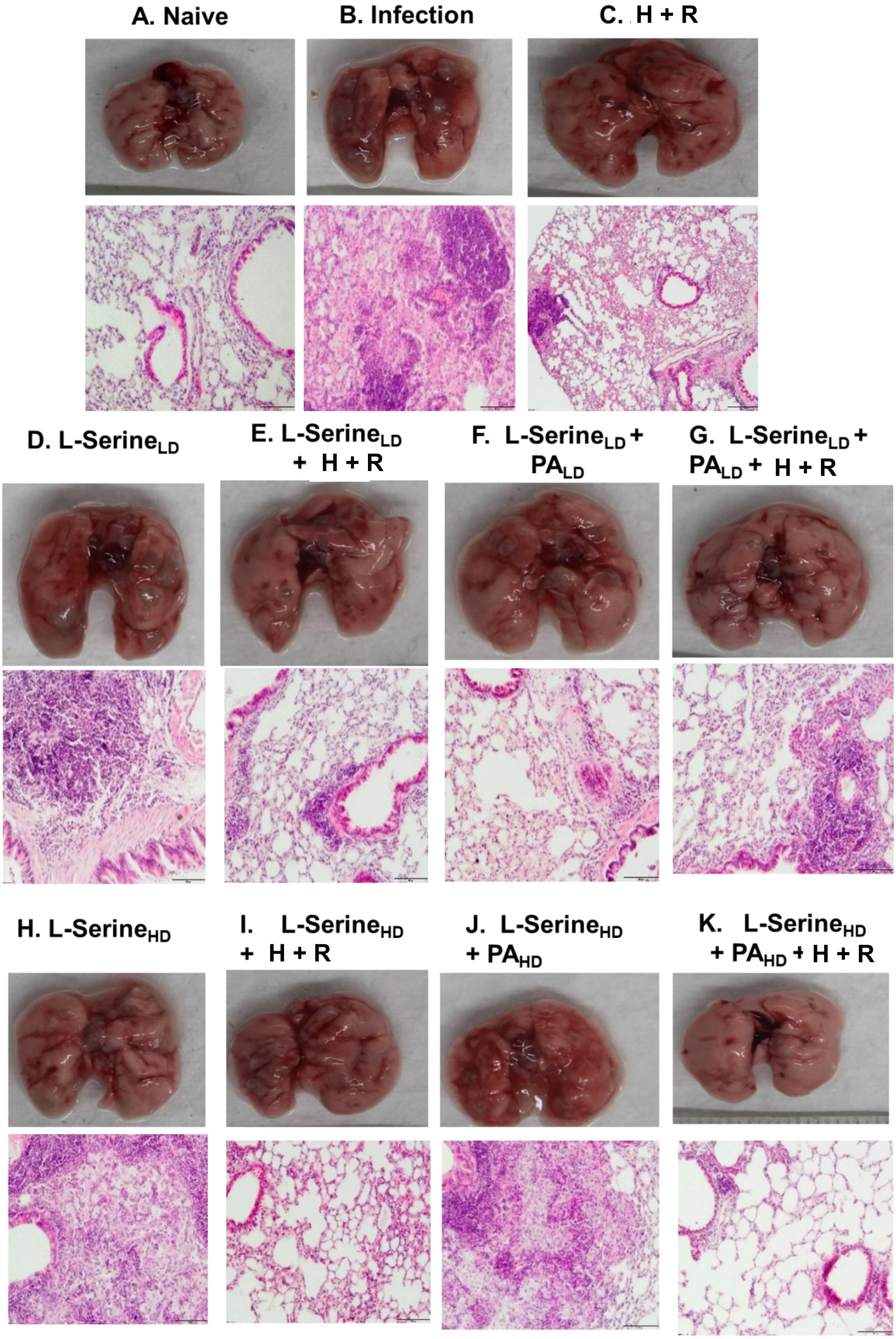

**Suppl. Figure 5**

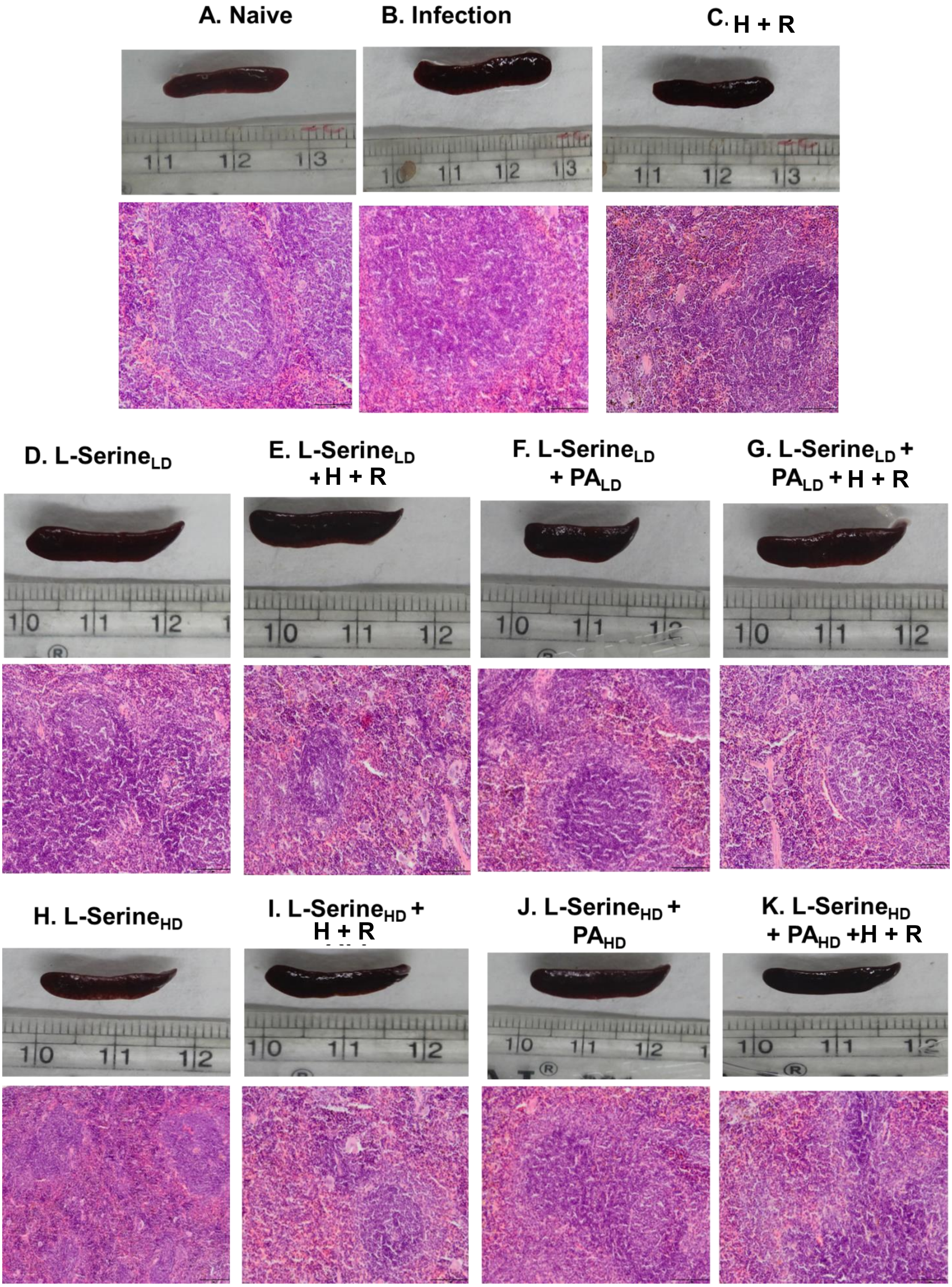

#### Supplementary Figures

**Suppl Figure 1. LS augment anti-tubercular efficacy of rifampicin against *Mycobacterium tuberculosis* $H_{37}$Rv.**

**(A)** Representative REMA plate images showing dose–response of rifampicin and L-Serine against *M.tb* $H_{37}$Rv. Wells shift from pink (metabolically active bacteria) to blue (growth inhibition) with increasing drug concentration. Approximate MICs are indicated: rifampicin ≈ 0.06 µM; L-Serine ≈ 50 mM. **(B)** Representative checkerboard REMA assessing combinations of rifampicin (columns; 0.003–0.06 µg/mL shown) and L-Serine (rows; 0.39–50 mM shown). Blue wells indicate inhibition; pink wells indicate growth. Data are representative of at least three independent experiments. **(C)** Representative REMA images for the MDR clinical isolate showing dose–response to moxifloxacin, rifampicin and L-Serine. **(D)** Checkerboard REMA for L-Serine + moxifloxacin demonstrates reduction of the effective moxifloxacin concentration to 0.125 µM in combination wells. **(E)** Checkerboard REMA for L-Serine + rifampicin shows enhancement of rifampicin activity at concentrations as low as 1.25 µM when combined with L-Serine. Images are representative of ≥3 independent experiments.

**Suppl Figure 2. LS and PCoA enhances the efficacy of Isoniazid and Rifampicin against MDR *M.tb***

**(A-B)** CFU count of MDR *M.tb* strain (SN/MC-90) treated with LS and P CoA alone or in combination at **(A)** 48Hrs and **(B)** 72Hrs. **(C-D)** CFU count of primary human $CD14^+$ monocytes treated with LS and H+R alone or in combination at **(C)** 48Hrs and **(D)** 72Hrs. Data are presented as mean ± SD from three independent experiments. Statistical significance was determined by one-way ANOVA followed by the appropriate post hoc test. **p** < 0.01; ***p*** < 0.001.

**Suppl. Figure 3. Lipid accumulation decreases in Foamy Macrophages upon LS treatment**

(A-B) Representative images of foamy macrophages from **(A)** healthy donor and **(B)** MDR-TB patients treated various lipid conditions oleic acid, naiveLDL, OxyLDL, and PA in combination with LS.

**Suppl Figure 4. L-Serine / Palmitic acid normalize lung morphology.**

Representative images of harvested lungs and hematoxylin and eosin (H&E) - stained lungs section micrographs show the impact of L-Serine and Palmitic Acid (PA), alone or in combination with H + R, on *Mycobacterium tuberculosis*–induced pulmonary pathology. Panels: **(A)** Naïve (uninfected); **(B)** Infection control (untreated); **(C)** H + R; **(D)** $LS^{LD}$ 600 mg/kg; **(E)** $LS^{LD}$ 600 mg/kg + H + R; **(F)** $LS^{LD}$ 600 mg/kg + $PA^{LD}$ 150 mg/kg; **(G)** $LS^{LD}$ 600 mg/kg + $PA^{LD}$ 150 mg/kg + H + R; **(H)** $LS^{HD}$ 1200 mg/kg; **(I)** $LS^{HD}$ 1200 mg/kg + H + R; **(J)** $LS^{HD}$ 1200 mg/kg + $PA^{HD}$ 300 mg/kg; **(K)** $LS^{HD}$ 1200 mg/kg + $PA^{HD}$ 300 mg/kg + H + R.

**Suppl. Figure 5. L-Serine / Palmitic acid mitigates splenomegaly in infected animals**

Spleen photographs and photographs and Hematoxylin & eosin–stained sections collected at the study end point show the impact of LS and PA, alone or in combination with H + R, on *M.tb*–induced pulmonary pathology. Panels: **(A)** Naïve (uninfected); **(B)** Infection control (untreated); **(C)** H + R; **(D)** $LS^{LD}$ 600 mg/kg; **(E)** $LS^{LD}$ 600 mg/kg + H + R; **(F)** $LS^{LD}$ 600 mg/kg + $PA^{LD}$ 150 mg/kg; **(G)** $LS^{LD}$ 600 mg/kg + $PA^{LD}$ 150 mg/kg + H + R; **(H)** $LS^{HD}$ 1200 mg/kg; **(I)** $LS^{HD}$ 1200 mg/kg + H + R; **(J)** $LS^{HD}$ 1200 mg/kg + $PA^{HD}$ 300 mg/kg; **(K)** $LS^{HD}$ 1200 mg/kg + $PA^{HD}$ 300 mg/kg + H + R.

**Supplementary Table 1: Checkerboard assay of Rifampicin with L-Serine against *M. tuberculosis H37Rv* and MDR Clinical Strain (SN/MC-90)**

Synergy is defined as FICI ≤ 0.5, 0.5 < FICI ≤ 1.0 as additive, 1.0 < FICI ≤ 4.0 as indifferent, and FICI > 4.0 as antagonism.

| ***M.tb* strain** | **Drug combination** | **MIC alone (L-Serine)** | **MIC alone (Drug)** | **MIC in combination (L-Serine)** | **MIC in combination (Drug)** | **FIC (L-Serine)** | **FIC (Drug)** | **FICI** | **Interaction** |
|---|---|---|---|---|---|---|---|---|---|
| **$H_{37}Rv$** | L-Serine + Rifampicin | 50 mM | 0.07 µM | 0.78 mM | 0.01 µM | 0.0156 | 0.14 | 0.16 | Synergistic |
| **MDR strain (SN/MC-90)** | L-Serine + Moxifloxacin | 400 mM | 0.62 µM | 6.25 mM | 0.31 µM | 0.0156 | 0.50 | 0.51 | Additive |
| **MDR strain (SN/MC-90)** | L-Serine + Rifampicin | 400 mM | 2.5 µM | 0.78 mM | 1.5 µM | 0.0019 | 0.60 | 0.60 | Additive |

**Supplementary Table 2. Histopathology scoring of lungs and spleen of infected animals supplemented with L-Serine and Palmitic acid under experimentation**

The table lists experimental groups (G1 – G11), a concise description of dominant histopathological findings in the lung, and an overall severity summary expressed as an approximate pathology score and qualitative category. Scoring method: five key histopathological parameters were evaluated semi-quantitatively for each animal - granulomatous foci, inflammatory infiltrate, necrosis, atelectasis/consolidation and fibrosis. Each parameter was scored using a 0 to 3 scale: 0 = absent (–), 1 = mild (+), 2 = moderate (++), 3 = marked/severe (+++). The approximate total pathology score shown in the table is the sum of the individual parameter scores (range ≈ 0–15). The qualitative severity categories are derived from the summed score and are intended as a concise guide: No lesion (Score = 0), Mild pathology (Score ≈ 1–5), Moderate pathology (Score ≈ 6–10) and Severe pathology (Score ≈ 11–15). Values given as a range (e.g., ≈10–12) reflect inter-animal variability within the group. Notes: “Isoniazid and Rifampicin” denotes standard anti-tubercular therapy; “PA” denotes palmitic acid.

| Group | Key Findings | Severity Summary |
|---|---|---|
| Naïve (Uninfected) | Normal pulmonary architecture without inflammation or fibrosis. | No lesion (Score: 0) |
| Infection (2 months) | Severe granulomatous inflammation (+3), PMN infiltration (+3), moderate necrosis (+2), moderate atelectasis (+2), moderate fibrosis (+2). | Severe pathology (Score ≈ 16–18) |
| Isoniazid and Rifampicin | Small granulomatous foci (+1), mild inflammatory infiltrate (+2), mild necrosis (+1), moderate atelectasis (+2), moderate fibrosis (+2). | Mild-to-moderate pathology (Score ≈ 9–10) |
| L-Serine (600 mg/kg) | Extensive granulomas (+3), marked inflammation (+3), moderate necrosis (+2), severe atelectasis (+3), moderate fibrosis (+2). | Severe pathology (Score ≈ 15–16) |
| L-Serine (600 mg/kg) + Palmitic Acid (150 mg/kg) | Moderate foci (+2), inflammation (+2), mild necrosis (+1), severe atelectasis (+3), mild fibrosis (+1). | Moderate pathology (Score ≈ 11–12) |
| L-Serine (600 mg/kg) + Isoniazid and Rifampicin | No granuloma, moderate inflammation (+2), no necrosis, moderate atelectasis (+2), mild fibrosis (+1). | Mild pathology (Score ≈ 6–7) |
| L-Serine (600 mg/kg) + Palmitic Acid (150 mg/kg) + Isoniazid and Rifampicin | Moderate foci (+2), mild inflammation (+1), no necrosis, mild atelectasis (+1), mild fibrosis (+1). | Mild pathology (Score ≈ 5–6) |
| L-Serine (1200 mg/kg) | Moderate granulomas (+2), severe inflammation (+3), moderate necrosis (+2), moderate atelectasis (+2), moderate fibrosis (+2). | Moderate-to-severe pathology (Score ≈ 14–15) |
| L-Serine (1200 mg/kg) + Palmitic Acid (300 mg/kg) | Mild granulomas (+1), moderate inflammation (+2), mild necrosis (+1), mild atelectasis (+1), mild fibrosis (+1). | Mild-to-moderate pathology (Score ≈ 7–8) |
| L-Serine (1200 mg/kg) + Isoniazid and Rifampicin | No granuloma, moderate inflammation (+2), no necrosis, normal lung (–), normal fibrosis (–). | Mild pathology (Score ≈ 3–4) |
| L-Serine (1200 mg/kg) + Palmitic Acid (300 mg/kg) + Isoniazid and Rifampicin | No granuloma, normal lung parenchyma (–) in all parameters. | No detectable pathology (Score: 0) |

**Supplementary Table 3. Primers used in this Study**

| S.No. | Genes | Primers |
|---|---|---|
| **1.** | SPHK1 | (F) 5’- CTTCACGCTGATGCTCACTG – 3’<br>(R) 5’ – GTTCACCACCTCGTGCATC – 3’ |
| **2.** | SPHK2 | (F) 5’ – GAGGAAGCTGTGAAGATGCCTG – 3’<br>(R) 5’ – GAGCAGTTGAGCAACAGGTCGA – 3’ |
| **3.** | SPTLC2 | (F) 5’ – ACGGAACGGGTACGTGAG – 3’<br>(R) 5’ – TTTGTGTAACATGATGGATCTGG – 3’ |
| **4.** | CERS1 | (F) 5’ – GTCACCCTGCAACCGTGCCAC – 3’<br>(R) 5’ – AGGTCGAAGACGACTGTCCACT – 3’ |
| **5.** | SMPD1 | (F) 5’ – CTCTTCCTCACTGACCTGCA -3’<br>(R) 5’ – CACCATATCAAAAGGGCCGC – 3’ |
| **6.** | Beta-actin | (F) 5’- AACTACCTTCAACTCCATCA -3’<br>(R) 5’- GAGCAATGATCTTGATCTTCA -3’ |

#### Annexure Table - Abbreviations Table used in this Study

| | |
|---|---|
| **WHO** | World Health Organization |
| ***M.tb*-$H_{37}$Rv** | *Mycobacterium tuberculosis* |
| **TB** | Tuberculosis |
| **MDR-PTB** | Multi-Drug-Resistant Pulmonary Tuberculosis |
| **XDR-TB** | Extensively-Drug Resistant Tuberculosis |
| **LS** | L-Serine |
| **PA** | Palmitic acid |
| **PCoA** | Palmitoyl CoA |
| **H** | Isoniazid |
| **R** | Rifampicin |
| **nLDL** | Naïve Low-Density Lipoprotein |
| **OxyLDL** | Oxidized Low Density Lipoprotein |
| **LPS** | Lipopolysaccharide |
| **$LS^{LD}$** | L-Serine Low Dose |
| **$LS^{HD}$** | L-Serine High Dose |
| **$PA^{LD}$** | Palmitic acid Low Dose |
| **$PA^{HD}$** | Palmitic acid High Dose |
| **NO** | Nitric Oxide |
| **$PCoA^{HD}$** | Palmitoyl CoA High Dose |
| **OA** | Oleic acid |
| **HC** | Healthy Control |
| **CFU $ml^{-1}$** | Colony Forming Units per mL |
| **TNF-α** | Tumor Necrosis Factor-alpha |
| **IFN-γ** | Interferon-gamma |
| **IL-6** | Interleukin-6 |
| **IL-10** | Interleukin-10 |
| **FICI** | Fractional Inhibitory Concentration Index |
| **REMA** | Resazurin Microtiter Assay |
| **HDT** | Host Directed Therapy |
| **ADC** | Albumin Dextrose Catalase |
| **OADC** | Oleic acid, Albumin, Dextrose, catalase |
| **7H9** | Middlebrook 7H9 Broth |
| **7H11** | Middlebrook 7H11 Agar |
| **SN/MC-90** | MDR *M.tb* strain Resistant against isoniazid and Rifampicin from Sarojini Naidu Medical College |
| **HRZE** | Isoniazid, Rifampicin, Pyrazinamide, Ethambutol |
| **BPaLM** | Bedaquiline, Pretomanid, Linezolid, Moxifloxacin |
| **LMICs** | Low- and Middle- Income Countries |
